# Restoring Parkin Function: An AAV Gene Therapy Approach for Early-Onset Parkinson’s Disease

**DOI:** 10.64898/2026.07.09.737487

**Authors:** Sambuddha Basu, Tyler G Demarest, Nicholas J. Gattone, Alexandra J Gilsrud, Brittany Wicks, Apeksha Khatiwada, Mohammad Nayal, Renee Gentzel, Daniel Cohen, Eric W. Kostuk, Derek P Narendra, Pedro Gonzalez-Alegre, Maria-Grazia Biferi, Elizabeth A Ramsburg

**Affiliations:** Spark Therapeutics, Philadelphia, PA, USA; Roche Innovation Center (RICP), Philadelphia, PA, USA; National Institutes of Health (NIH), National Institute of Neurological Disorders and Stroke (NINDS), Bethesda, MD, USA

**Keywords:** Early Onset Parkinson’s Disease (EOPD), Parkin, mitophagy, phosphorylated ubiquitin at serine 65 (pUb^Ser65^), AAV-gene therapy, gene replacement therapy

## Abstract

**Background:** Biallelic loss-of-function mutations in *PRKN* gene (encoding Parkin protein) cause early-onset Parkinson’s disease (EOPD). Parkin is a crucial component of PINK1-Parkin pathway, which marks damaged mitochondria for degradation via mitophagy. Without functional Parkin, damaged mitochondria accumulate, causing oxidative stress and neurodegeneration.

**Objective:** Investigate Parkin gene replacement via AAV gene therapy as a potential treatment for Parkin-dependent EOPD.

**Methods:** We initially validated phosphorylated ubiquitin Ser65 (pUb^Ser65^) as an indicator of Parkin-mediated mitophagy initiation. We evaluated AAV-mediated *PRKN* replacement (hereafter, AAV-Parkin) in a Parkin knockout neuroblastoma cell line (SH-SY5Y cells) and feasibility of delivery in mouse and rat models.

**Results:** Our research showed pUb^Ser65^ signal was reduced in Parkin-KO SH-SY5Y cells when compared to wild-type cells after mitochondrial stress, indicating deficiency in initiation of mitophagy. AAV-mediated human *PRKN* gene replacement successfully restored these pUb^Ser65^ levels in knockout cells. We saw restoration in patient-derived fibroblasts following AAV-Parkin overexpression. We developed a translatable gene therapy approach using rodents. We demonstrated the feasibility of delivering AAV-Parkin directly into the substantia nigra (SN) of wild-type rats. Using an AAV1 capsid with Ef1a promoter, we achieved dose-dependent Parkin expression and identified a well-tolerated dose. We also evaluated multiple promoters in a proprietary Spark100 capsid, finding Ef1a and Synapsin1 (Syn1) were most effective for transducing dopaminergic neurons in the SN of mice without causing adverse effects. These findings established a well-tolerated vector dose and an optimal capsid-promoter combination.

**Conclusions:** Our results support the potential of AAV-Parkin gene therapy as a disease-modifying approach for Parkin-deficient EOPD.

**Graphical Abstract:** 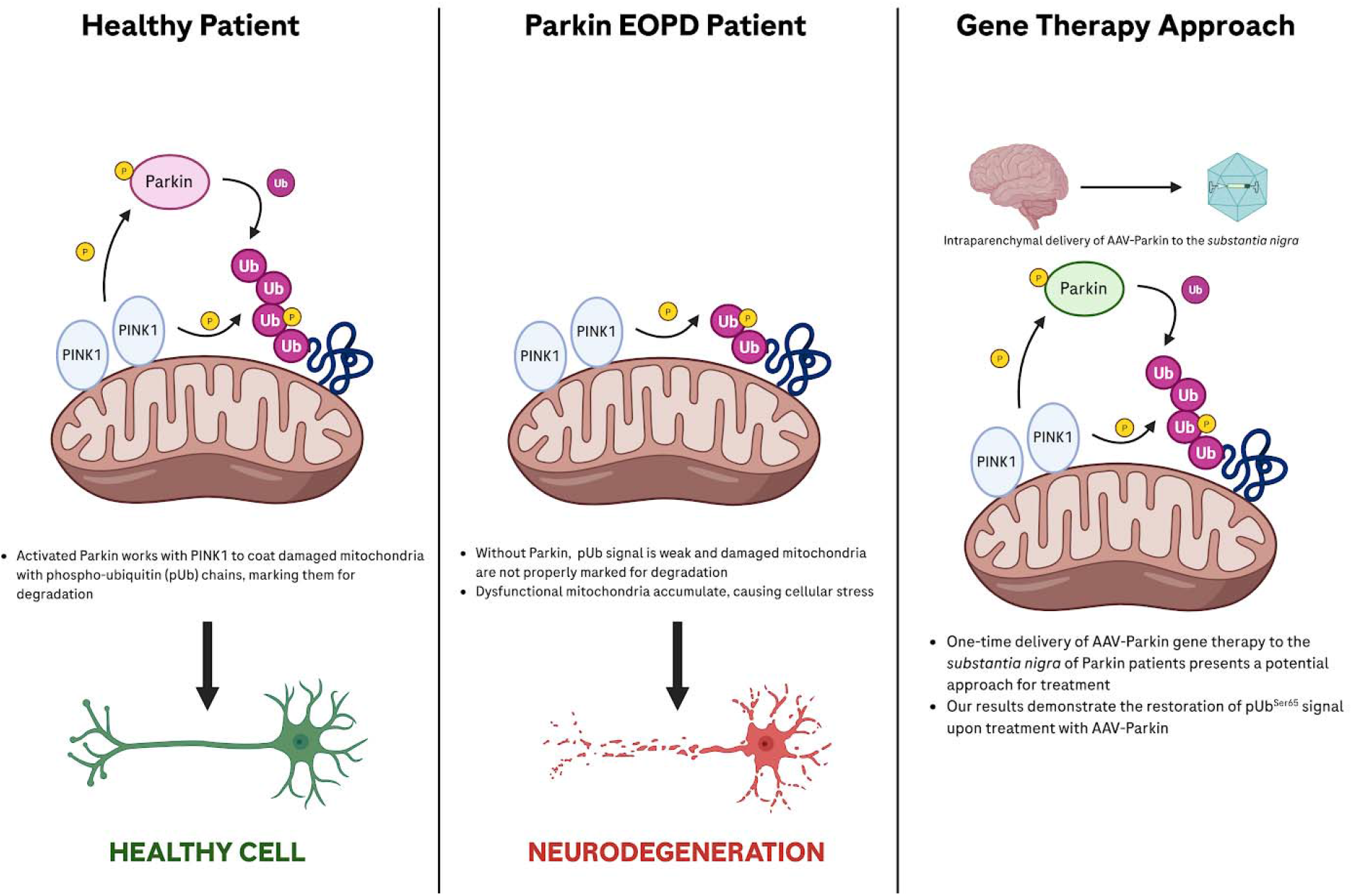

## Introduction

Parkinson’s disease (PD) is a progressive neurodegenerative disorder characterized by motor symptoms including bradykinesia, rigidity, tremor, and postural instability caused by degeneration of dopaminergic neurons of substantia nigra pars compacta (SN/SNpc)^1^. While majority of PD cases are classified as idiopathic and late-onset, a subset of patients exhibits early-onset Parkinson’s disease (EOPD), defined as the onset of symptoms before the age of 50^2^. EOPD is clinically and genetically distinct from late-onset PD, with a higher likelihood of genetic contributions playing a central role in its pathogenesis.

The commonly mutated gene in EOPD is *PRKN*, which in some populations is found in approximately 2.6%-14.9% of EOPD cases^4, 5, 6^. Epidemiological studies provide insight into the prevalence of *PRKN* mutations. Two recent studies estimated the genetic prevalence of PRKN-PD to be approximately 13 per 100,000 ^7, 8^. The *PRKN* gene, which encodes E3 ubiquitin ligase Parkin, plays a crucial role in maintaining cellular homeostasis through the regulation of protein degradation and mitochondrial function^4, 9^. Loss-of-function mutations in the *PRKN* gene lead to impaired mitophagy, an essential process for the removal of damaged mitochondria, which contributes to the accumulation of defective proteins and cellular stress, ultimately resulting in neurodegeneration and EOPD^10^. The PINK1-Parkin pathway is a crucial process for clearing damaged mitochondria through mitophagy. When mitochondria are stressed, PINK1 accumulates on their surface and phosphorylates ubiquitin at Serine 65 (pUb^Ser65^), a key signal for the initiation of mitophagy. Binding phosphorylated ubiquitin is the first event required for Parkin activation. PINK1 then directly phosphorylates Parkin itself, providing the second signal required for activation of Parkin’s E3 ligase activity. Parkin, once activated, ubiquitinates proteins on the mitochondrial surface, providing an additional substrate for PINK1. This combined action greatly amplifies the damage signal, recruiting more Parkin to the damaged mitochondria and initiating the full mitophagy process^11^. The accumulation of pUb^Ser65^ requires PINK1 but is enhanced by endogenous Parkin.

Disease-causing mutations in *PRKN* range from single base pair substitutions, small deletions, and splice site mutations to deletions and duplications spanning many nucleotides^10^. *PRKN* mutations lead to a loss of Parkin function. For missense mutations, loss of Parkin function may be due to decreased catalytic activity and/or destabilization of Parkin leading to insolubility or rapid proteasomal degradation of mutant Parkin. Gene therapy using adeno-associated virus (AAV) vectors offers a promising strategy for treating *PRKN*-associated EOPD by delivering functional *PRKN* gene (and Parkin protein) to the SN^12^. AAV vectors are particularly well-suited for gene therapy due to their safety, ability to transduce both neuronal and non-neuronal cells, and long-term expression in the target tissue^12^. Direct delivery of *PRKN* gene (and Parkin protein) to SN can potentially restore its function in dopaminergic neurons where endogenous Parkin function is lost, enhancing mitochondrial quality control and promoting the clearance of damaged mitochondria through mitophagy^13^. This approach could mitigate neurodegeneration of dopaminergic neurons and the development of PD symptoms. Gene-therapy approaches delivering AAV vector payload directly to SN and dopaminergic neurons have not only been well-established in clinics but also deemed safe and a reasonable translatable approach^14,15^.

In this manuscript, we evaluate AAV gene therapy for *PRKN*-associated EOPD. Our findings highlight increased pUb^Ser65^ as an indicator of Parkin activation when mitochondria are stressed. We further show that AAV-Parkin gene therapy can restore this activation. Building on our in-vitro findings, we next optimized promoters to ensure the efficient and targeted delivery of functional Parkin to dopaminergic neurons in the SN of wild-type rodents. Our foundational hypothesis is that restoring at least 50% of the endogenous functional Parkin protein expression in dopaminergic neurons will be therapeutically beneficial. This is supported by human genetics, which has established that Parkinson’s disease (PD) only develops in individuals with biallelic loss of function in the *PRKN* gene^8^. This clinical observation provides a strong rationale for our gene therapy approach, as restoring even a single functional copy of the gene should be sufficient to prevent disease progression. We demonstrate *in vivo* feasibility and dose-finding studies to achieve targeted 50% endogenous Parkin expression in the SN of rats. Our work proves the foundational concept and outlines a clear path for translating AAV-Parkin gene therapy into a potentially clinically viable, disease-modifying treatment for *PRKN*-deficient EOPD.

## Methods

Detailed descriptions of all materials and methods can be found in the Supplementary material. These include assessment of WT and Parkin knock out SH-SY5Y cells, patient derived fibroblasts from healthy controls and Parkin-PD patients for initiation of mitophagy signal, tolerability assessment of AAV-Parkin gene therapy and promoter optimization.

## Data availability

All data are provided in the main text or supplementary section, with no data deposited in external repositories. All associated data presented here are available on reasonable request from the corresponding authors (S.B and M.G.B).

## Results

### AAV-Parkin rescues pUb^Ser65^ signal and mitophagy initiation following mitochondrial stress in Parkin-deficient cells

To determine a cellular phenotype consistent with Parkin deficiency, we evaluated a commercially available Parkin knockout (KO) SH-SY5Y neuroblastoma cell line. Initial experiments revealed no differences in mitophagy and ubiquitin proteasome system endpoints under basal conditions between WT and Parkin KO cells treated with DMSO (Fig 1A). We then evaluated the addition of mitochondrial stress to mimic the age-related accumulation of damaged mitochondria that contributes to dopaminergic neuron dysfunction as observed in *PRKN*-EOPD patients. The loss of mitochondrial membrane potential limits cellular energy production in the form of ATP and initiates the removal of uncoupled mitochondria through both Parkin-dependent and Parkin-independent mitophagy pathways. Therefore, we evaluated the pharmacological proton ionophore to collapse mitochondrial membrane potential to mimic age-related mitochondrial impairments using carbonyl cyanide 4-(trifluoromethoxy)phenylhydrazone (FCCP) as a stimulus to initiate Parkin-dependent mitophagy. We identified a Parkin-mediated significant increase in pUb^Ser65^ in WT cells only, not in the Parkin KO neuroblastoma cells. This is consistent with literature demonstrating that pUb^Ser65^ production by PINK1 can be amplified by Parkin^15^.

**Figure 1.**
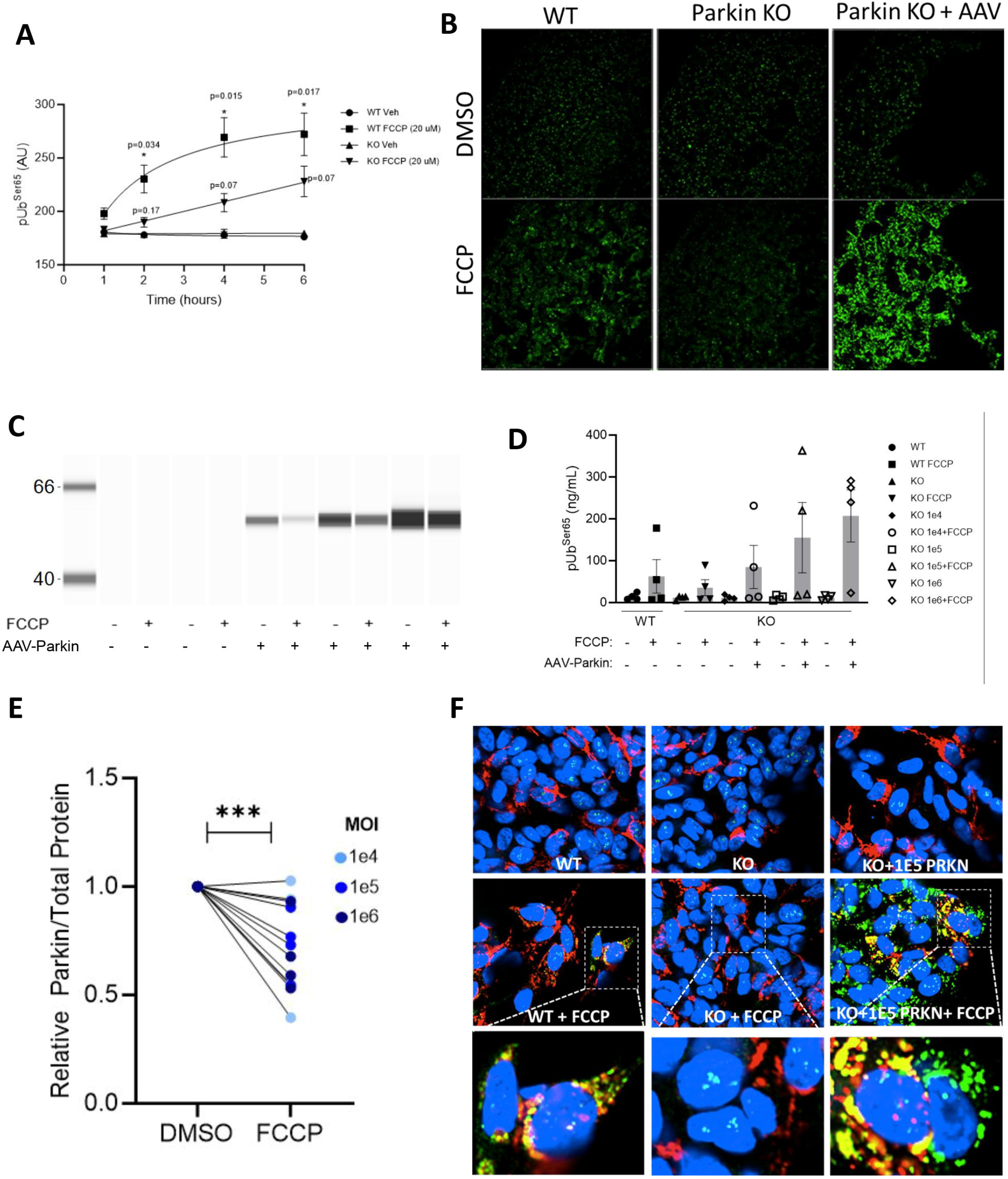
AAV-Parkin (human) rescues pUb^Ser65^ deficiency following mitochondrial stress in Parkin Knock out (KO) SH-SY5Y Neuroblastoma cells. A, B) Time-course identification of pUb^Ser65^ deficiency in Parkin KO cell lysates following vehicle or FCCP (20 µM) by dot blot and confirmed by immunocytochemistry staining. *p<0.05, 2-way ANOVA vs. vehicle treatment, RM-2-Way ANOVA with Tukey’s post hoc test. C) AAV1-Ef1a-Parkin (human) transduction of Parkin KO cells reveals MOI-dependent expression of Parkin and D) pUb^Ser65^ by ELISA E) Parkin protein is reduced following FCCP treatment reflecting Parkin degradation following mitochondrial stress in KO cells overexpressing Parkin by AAV-Parkin. ***p=0.0005, Student’s t-test DMSO vs. FCCP in KO cells expressing Parkin by AAV-Parkin. n=12 independent lysates.. F) Parkin co-localizes with mitochondria-RFP following FCCP treatment.

An initial time-course evaluation of pUb^Ser65^ by dot-blot analysis demonstrated induction of pUb^Ser65^ in WT cells (starting at 2hrs and peaking at 4hrs) when challenged with 20uM FCCP, compared to DMSO treated WT cells, indicating a normal induction of mitophagy (Fig 1A, WT vs. WT+FCCP, p<0.05 at 2hrs, 6hrs and 8hrs, Supplementary Fig 1A). However, in Parkin KO cells, this showed a non-significant change in pUb^Ser65^ signal following FCCP treatment at all time-points compared to control Parkin KO cells with DMSO (Fig 1A, KO vs. KO+FCCP, p>0.05 for all time points, Supplementary Fig 1A). This result was confirmed qualitatively through pUb^Ser65^ immunofluorescence staining (Fig 1B, panel 1 and 2 – top and bottom). After endpoint identification of Parkin-dependent pUb^Ser65^ production, we interrogated whether deficiency in pUb^Ser65^ production could be restored by introducing a functional copy of the *PRKN* gene using gene therapy.

WT and Parkin KO cells were transduced with 3 Multiplicity of infection (MOI) of AAV1-Parkin (human) driven by the ubiquitous promoter Ef1a. Following 72 hours of transduction, cells were treated with FCCP or DMSO (vehicle control) and harvested for Parkin protein expression and pUb^Ser65^ determination by ELISA. We observed MOI-dependent increase in Parkin protein expression (Fig 1C), and similar dose-dependent trends for pUb^Ser65^ upregulation following FCCP, but not vehicle treatment, indicating AAV-Parkin was able to upregulate pUb^Ser65^ (Fig 1D). This amplification of pUb^Ser65^ signal was visually confirmed through pUb^Ser65^ immunofluorescence staining in Parkin KO cells treated with optimal MOI of AAV1-Ef1a-Parkin (human) and challenged with FCCP (Fig 1B, panel 3 – top and bottom). We then investigated whether the damaged mitochondria of Parkin KO cells were able to be degraded by mitophagy after AAV-Parkin treatment. Notably, we observed a statistically significant reduction of Parkin protein in the cells treated with FCCP compared to the DMSO control (Fig 1E, p=0.0005), indicating the gene therapy delivered Parkin was likely binding to dysfunctional mitochondria and being degraded alongside mitochondrial cargo via mitophagy. To further investigate the mitochondrial localization of AAV-Parkin, we performed co-staining experiments with mitochondrial-tagged red fluorescence protein (mito-RFP) and human Parkin protein (transduced by AAV1-Ef1a-Parkin) in Parkin KO cells treated with DMSO vs. FCCP. We observed absence of Parkin staining in untreated and non-transduced KO cells, and a cytoplasmic staining pattern in AAV-Parkin (green) DMSO treated cells. Importantly, we observed mitochondrial co-localization with AAV-delivered Parkin protein following FCCP treatment (colocalization of mito-RFP and green signal of Parkin, Fig 1F). Together, these experiments demonstrate pUb^Ser65^ amplification is deficient in Parkin KO neuroblastoma cells following mitochondrial damage, but can be restored following AAV-Parkin treatment.

### AAV-Parkin restores pUb^Ser65^ amplification signal in patient-derived fibroblasts from EOPD (PRKN-PD) following mitochondrial stress

While the initial results suggest a promising approach for the restoration of Parkin function in Parkin KO SH-SY5Y cells, patient-relevant *PRKN* gene mutations contributing to EOPD are diverse. This genetic complexity may not be fully recapitulated in a KO cell system. Therefore, we evaluated patient-derived fibroblasts containing distinct *PRKN* mutations (n=8-9), or fibroblasts from healthy control non-Parkin mutation carriers (n=4) (Table 1). All patient lines contained biallelic mutations in either a recessive or compound heterozygous state. To study how Parkin affects initiation of mitophagy, we first had to find an efficient way to transduce patient-derived fibroblasts. We compared different AAV (adeno-associated virus) capsids and found that AAV2 was the most effective. This vector, which initially used a CAG promoter to express a GFP (green fluorescent protein) transgene, proved successful (Supplementary Fig 2A). With this optimized method, we created a standardized procedure to deliver the human *PRKN* gene into the fibroblasts using the same AAV2-CAG vector. For the next round of experiments, we treated these cells with valinomycin, a mitochondrial uncoupler, to observe how the newly introduced Parkin protein would respond to mitochondrial stress, as shown in the schematic image (Fig 2A).

**Figure 2:**
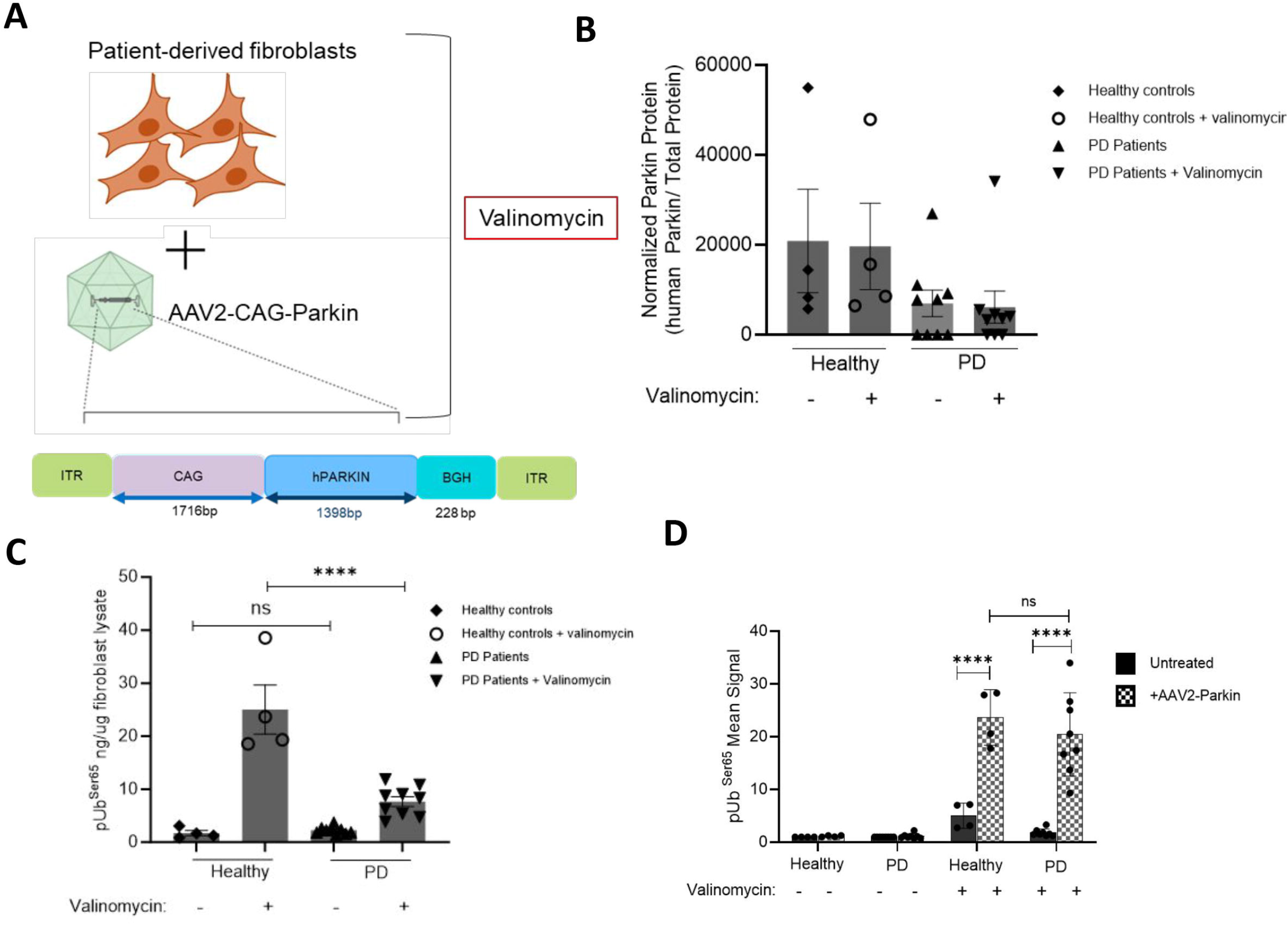
AAV-Parkin restores pUb^Ser65^ deficiency following mitochondrial stress in PD patient-derived fibroblast lines. (A) Graphical representation showing fibroblast lines from PD patients and controls from non-PD first degree relatives were treated with AAV2-CAG-Parkin (human) and stressed with valinomycin B) Quantified Parkin assessment by capillary electrophoresis shows no change between PD patients and controls from non-PD, and also after stress with Valinomycin. (C) pUb^Ser65^ by ELISA showing deficit of pUb^Ser65^ in fibroblasts from PD patients when compared to controls from non-PD. ****P<0.0001, Ordinary one-way ANOVA Tukey’s multiple comparisons test. (D) Quantified mean signal from pUb^Ser65^ ICC staining demonstrating restoration of pUb^Ser65^ signal comparable to WT levels when cells are treated with AAV2-CAG-Parkin (human) and stressed with valinomycin. ****P<0.0001 2 way ANOVA Tukey’s multiple comparisons test. Selected statistical comparisons are shown on the graph.

Basal levels of Parkin protein expression were similar with and without valinomycin treatment (Fig 2B). However, a significant reduction in pUb^Ser65^ signal was observed in Parkin mutation carriers compared to healthy controls when challenged with valinomycin, when measured by both ELISA and immunocytochemistry (Fig 2C, p<0.0001, Supplementary Fig 2B). We did not observe any difference in the basal level of pUb^Ser65^ between Controls and PD-patient cells. Following treatment with valinomycin, fibroblasts from *PRKN*-PD patients showed decreased production of pUb^Ser65^, consistent with absent Parkin activity in these cells.

For the next set of experiments, we aimed at rescuing the pUb^Ser65^ production defect in *PRKN*-PD patients by transducing the fibroblasts with AAV2-CAG-Parkin (human, MOI:1e4 particles/condition) (Fig 2D). We demonstrated by immunocytochemistry that replacing functional Parkin with AAV2-CAG-Parkin (human) rescued deficient pUb^Ser65^ amplification following valinomycin treatment in all *PRKN*-PD patient lines irrespective of the Parkin mutations and endogenous expression. pUb^Ser65^ in *PRKN*-PD patient cells after rescue was comparable to the upregulation of pUb^Ser65^ signal observed in healthy control lines (Fig 2D, Supplementary Fig 2B, Supplementary Fig 2C, p<0.0001). Individual data points from each fibroblast line post transduction with AAV2-CAG-Parkin (human) are shown in Supplementary Fig 2C. Together, these results demonstrate AAV-Parkin gene therapy restores Parkin activity in Parkin-mutant cells, irrespective of mutation.

### Intraparenchymal substantia nigra (SN) injection of AAV-Parkin in rats is feasible and well tolerated

We next investigated the feasibility and tolerability of delivering human Parkin by AAV-gene therapy by stereotaxic intraparenchymal (IPa) injection in the SN. To evaluate this, we administered 3 doses (3e8, 3e9 and 3e10 vg/SN) of AAV1-Ef1a-Parkin (human) to 2-month-old rats. The animals’ body weights remained comparable after treatment (Fig 3A). We observed a dose-dependent increase in immunostaining of human Parkin protein (red), which co-localized with the dopaminergic neuron marker tyrosine hydroxylase (TH in green; Fig 3B), indicating successful delivery to SN. Using capillary electrophoresis western blotting, we quantified Parkin protein compared to naïve and diluent injected rat SN samples and observed increasing dose-dependent expression of human Parkin protein (Fig 3C, p<0.05, Supplementary Fig 3A). To further understand the amount of human Parkin protein delivered in relation to endogenous levels of rat Parkin expression, we utilized an antibody that detects both human and rat Parkin protein. We observed that the 3e8vg/SN dose of AAV-Parkin resulted in Parkin expression ranging from 35 - 260% (mean=135.25%) compared to endogenous rat Parkin, which was slightly above the desired range of 50-100%. These data suggest that we successfully identified a dose that could potentially replenish Parkin levels to a functional range (Fig 3D, Supplementary Fig 3B).

**Figure 3.**
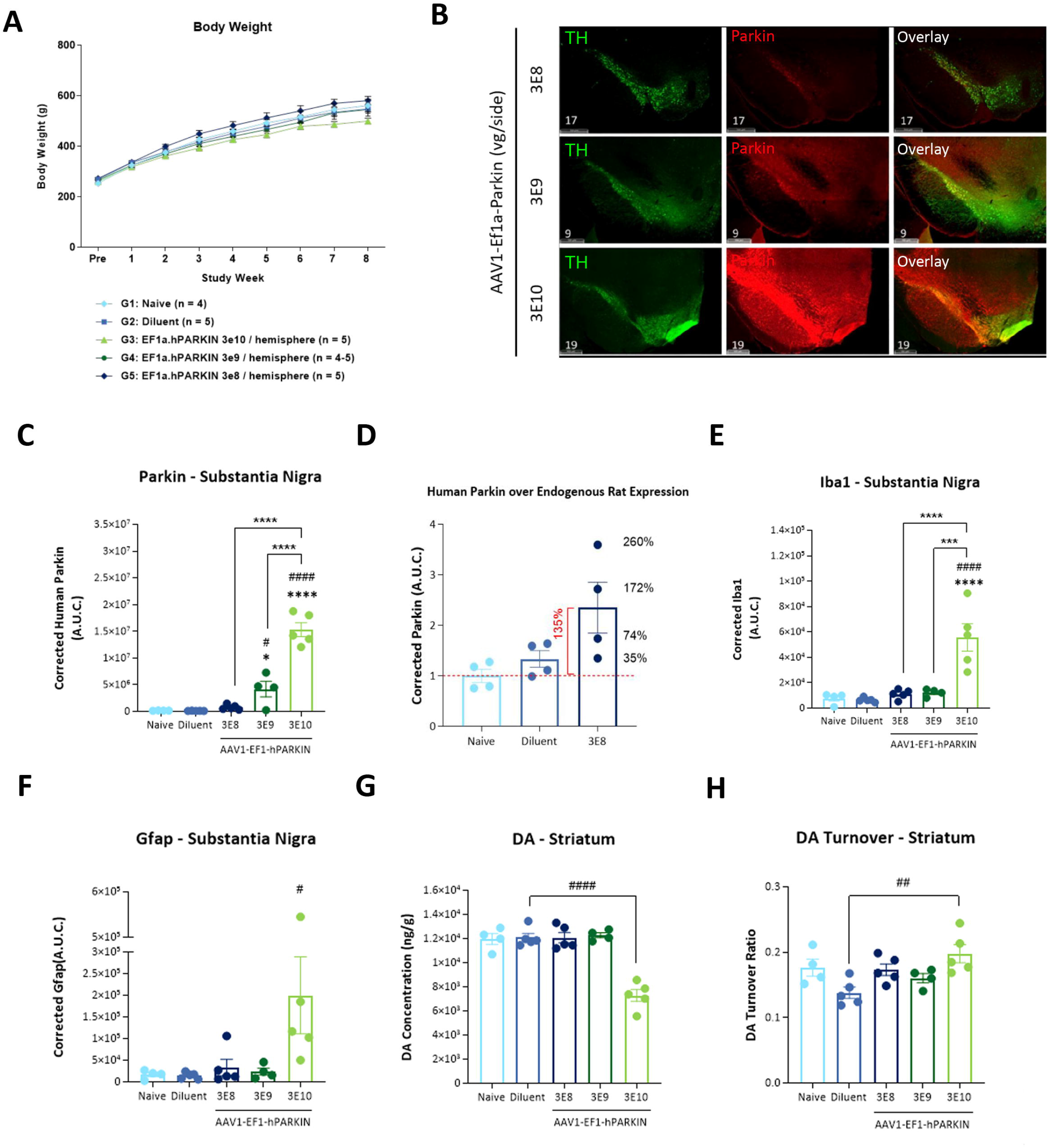
Dose-dependent Parkin (human) expression following stereotaxic delivery of AAV-Ef1a-Parkin (human) in Rat substantia nigra pars compacta (SNpc). (A) Rats bilaterally injected in the SN using stereotaxic surgical procedures showed increased body weight over time (B, C) Brain sections stained for TH (dopaminergic neuron marker, green) and human Parkin protein (red) shows dose-dependent increase of Parkin expression in SN and colocalization with dopaminergic neuron, which is also confirmed by capillary electrophoresis based biochemical assessment of human specific Parkin. 1-way ANOVA with Tukey’s post hoc analysis performed showing comparisons as Naïve vs 3E9, *p= 0.0412; Naïve vs 3E10, ****p 0.0001; Diluent vs. 3E9, ^#^p = 0.0276; Diluent vs 3E10, ^####^p<0.0001; 3E8 vs 3E10, ****p<0.0001; 3E9 vs 3E10, ****p<0.0001; (D) Human Parkin and total Parkin (human and mouse) quantified by capillary electrophoresis in the 3e8 vg/SN cohort to determine therapeutic levels of Parkin delivered with a goal of reaching at least 50% Parkin expression. (E, F) To assess tolerability, reactive markers for astrocytes (GFAP, E) and microglia (Iba1, F) were quantified by capillary electrophoresis. 1-way ANOVA with Tukey’s post hoc analysis performed showing comparisons of GFAP-Diluent vs 3E10, ^#^p=0.0471; IBA1-Naïve vs 3E10, ****p 0.0001; Diluent vs 3E10, ^####^p<0.0001; 3E8 vs 3E10, ****p<0.0001; 3E9 vs 3E10, ***p=0.0003; (G, H) Striatal dopamine and dopamine turnover quantified by HPLC showed potential disruption of neurotransmitter synthesis at the highest dose of 3e10vg/SN only. 1-way ANOVA with Dunnett’s post hoc analysis performed showing comparisons of Dopamine: Diluent vs 3E10, ^####^p<0.0001 and Dopamine turnover: Diluent vs 3E10, ^###^p=0.0026

To assess the tolerability of Parkin protein expression, we measured glial markers Gfap and Iba1 to screen for astrocyte and microglial activation, respectively. We observed significant upregulations of Gfap (Fig 3E, Diluent vs. 3e10 vg, #p=0.0471, Supplementary Fig 3C) and Iba1 (Fig 3F ####p<0.0001, Supplementary Fig 3D) at 3e10vg/SN only. Our study lacked control for the AAV vector itself at the 3e10 vg/SN dose, meaning we cannot definitively separate the vector’s effect from that of Parkin protein overexpression. However, the rise in inflammatory markers at this high dose potentially suggests that an excess of Parkin protein in the SN might be activating a neuroinflammatory response. As decreased striatal dopamine is a hallmark of nigrostriatal degeneration associated with PD, we quantified striatal dopamine and metabolites by HPLC. We observed no difference in dopamine or dopamine turnover ([DOPAC+HVA]/Dopamine) at 3e8 or 3e9 vg/SN compared to the Diluent group. However, consistent with the potentially adverse glial activation observed, we observed significant reduction in striatal dopamine (Fig 3G p<0.0001) and impaired dopamine turnover (Fig 3H, p=0.0026) at the 3e10 vg/SN dose. Altogether, the results demonstrate that functional replacement of Parkin protein is feasible by direct SN IPa injection of AAV-Parkin starting at a dose of 3e8/SN and is well-tolerated at doses below 3e10 vg/SN, giving a wide therapeutic window for *PRKN* gene replacement.

### Ef1a and Syn1 promoters drive optimal Parkin expression in AAV vectors in the SN of wild-type mice

In the previous tolerability assessment, we utilized AAV1 serotype and a ubiquitous promoter (Ef1a) to drive the expression of the Parkin protein in dopaminergic neurons. We sought to enhance the specificity of Parkin expression to the dopaminergic neurons of SN by evaluating regulatory promoter sequences driving Parkin expression in a Spark proprietary capsid Spark100. Six AAV constructs expressing full length human Parkin driven by different promoters and a diluent control were delivered by intraparenchymal injection to the SN of wild-type mice at 1e10vg/SN. For comparability to prior experiments, we utilized the same AAV1-Ef1a-Parkin (human) construct injected in rats (Fig 3), and Spark100 vectors with 2 ubiquitous promoters (Ef1a or CAG), one neuron-specific promoter (human Synapsin; hSyn1), and 2 dopaminergic neuron-specific promoters (Tyrosine hydroxylase: TH – full length or mini) (Fig 4A)^17^. 8 weeks following AAV injection, mice were sacrificed and one hemibrain was collected for immunohistochemistry and the other for biochemical analysis. Parkin protein was assessed biochemically by capillary electrophoresis western blot. We observed the greatest Parkin expression in the SN with the ubiquitous CAG promoter, followed by Ef1a, TH-full length, and hSyn1, with a lack of expression by the TH-mini promoter respectively (Fig 4B, Supplementary Fig 4A). To further assess successful dopaminergic neuron-specific expression, we immunostained brain sections. These showed colocalization of human Parkin protein (red) with TH positive dopaminergic neurons (green), supporting the biochemical assessment of Parkin (Fig 4C). Notably, we observed that while the TH-full promoter drives Parkin expression, there appeared to be a disruption in TH immunoreactivity within the SN. To assess any potential impact of dopaminergic neuron viability, we evaluated dopaminergic neuron marker TH by capillary-immunoblot and observed no statistically significant differences (Fig 4D, Supplementary Fig 4B). The Parkin protein expression in the striatum corroborated the observations in SN (Fig 4E). It is well reported in the literature that loss of TH positive dopaminergic fibers projecting in the striatum often precedes loss of TH cell bodies in the SN. Biochemical assessment of TH in striatum, showed a statistically meaningful lowering in Spark100-CAG-Parkin (human) and Spark100-TH (full)-Parkin (human) groups compared to Spark100-hSyn1-Parkin (human) group (Fig 4F). These data strengthened the trend observed in TH levels from SN and affirmed that hSyn1 and Ef1a were the most optimal promoters in the Spark100 capsid for targeting the dopaminergic neurons.

**Figure 4.**
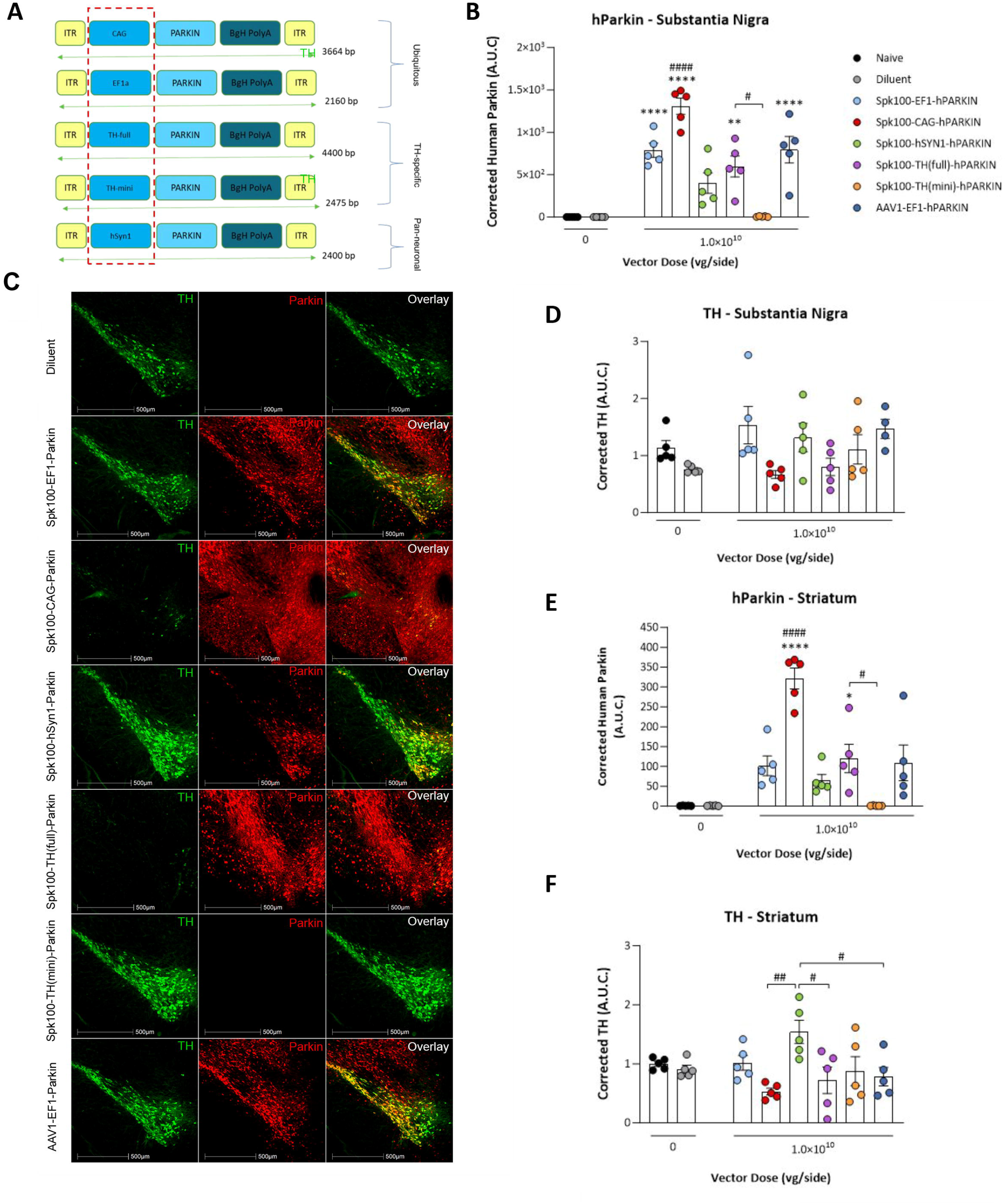
Selection of Optimal Promoter sequence for expressing Parkin(human) Expression in dopaminergic neurons of mouse SN. (A) Schematic of vector constructs expressing Parkin (human) under different promoters. Dotted red box indicates variable promoter regions used for the constructs. (B) The normalized expression of Parkin quantified by capillary electrophoresis (human) from different promoters in SN were plotted as individual values (symbols) and group means ± SEM (bars) and analyzed by a one-way ANOVA followed by Tukey’s post hoc test. Statistically significant pairwise comparisons between the controls (naïve or diluent) and the experimental groups were reported as follows: *p<0.5, and ***p<0.0001. Statistically significant pairwise comparisons between experimental groups were reported as follows: ^#^p<0.05, ^##^p<0.01, and ^####^p<0.0001 (C) Representative images of TH (green) and Parkin (human, red) overlay staining from animals injected with the diluent or dosed with 1.0×10^10^ vg/SN of Spark100-Ef1a-Parkin, Spark100-CAG-Parkin, Spark100-hSyn1-Parkin, Spark100-TH (full)-Parkin, Spark100-TH (mini)-Parkin, or AAV1-Ef1a-Parkin. (D) Normalized TH expression quantified by capillary electrophoresis in SN tissues following overexpression of Parkin (human) from different promoters at 1e10vg/SN. (E, F) Normalized expression of Parkin (human) and TH quantified by capillary electrophoresis from striatum tissues following overexpression of Parkin (human) from different promoters at 1e10vg/SN. Expression Values plotted as individual values (symbols) and group means ± SEM (bars) and analyzed by a one-way ANOVA followed by Tukey’s post hoc test. Statistically significant pairwise comparisons between the controls (naïve or diluent) and the experimental groups were reported as follows: p<0.01, and p<0.0001. Statistically significant pairwise comparisons between experimental groups were reported as follows: ^#^p<0.05, and ^####^p<0.0001 SNpc = substantia nigra pars compacta; TH = tyrosine hydroxylase; vg = vector genome.

Based on the prior observation from Fig 3E and Fig 3F, that high Parkin expression may result in adverse glial activation, we sought to determine whether there were promoter and expression-mediated alterations in astrocyte (GFAP) and microglial (Iba1) immunoreactivity in this study. Indeed, we observed that the ubiquitous CAG promoter driving the highest levels of Parkin protein expression in the SN also resulted in significant upregulation in Iba1 (Fig 5A p<0.0001, Supplementary Fig 5A) and GFAP (Fig 5B p<0.0001, Supplementary Fig 5B). Consistently, we observed reduced TH expression in SN and Striatum following Parkin expression from the ubiquitous CAG promoter. Importantly, all other constructs tested did not display any significant alterations in glial immunoreactivity (representative images Fig 5C). These results confirm that the dopaminergic-specific expression of Parkin is feasible and well-tolerated but suggest the importance of careful dose-finding studies to avoid toxic overexpression of Parkin in midbrain dopaminergic neurons.

**Figure 5.**
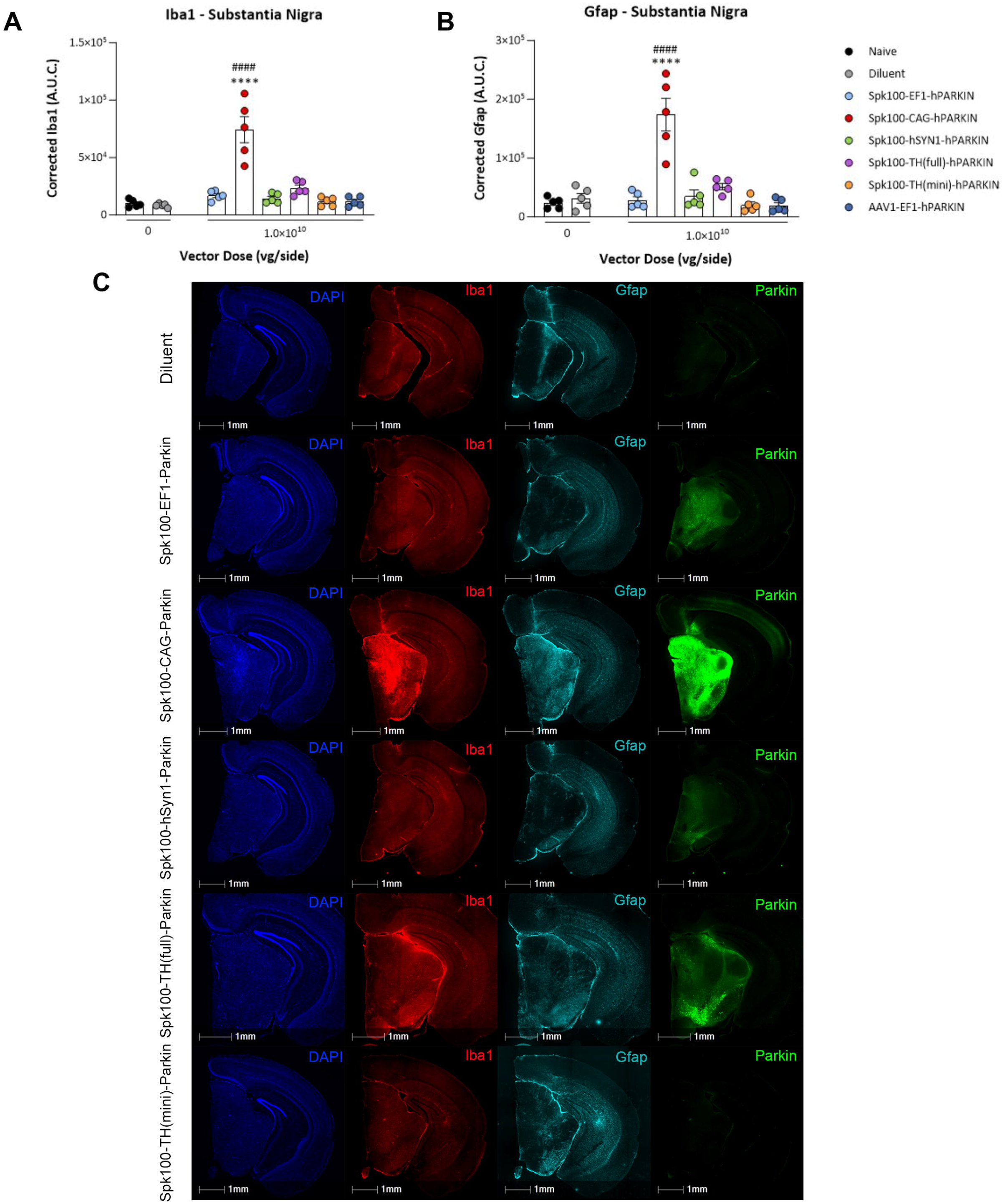
Evaluation of gliosis markers Iba1 and GFAP in mouse SN Tissues following overexpression of Parkin (human) from different promoters. (A, B) Normalized Iba1 and GFAP expression quantified by capillary electrophoresis in SN tissues following overexpression of Parkin (human) from different promoters were plotted as individual values (symbols) and group means ± SEM (bars) and analyzed by a one-way ANOVA followed by Tukey’s post hoc test. (C) Representative images of DAPI (blue), Iba11 (red), GFAP (cyan), and human Parkin (green) staining from animals injected with the diluent or dosed with 1e10 vg/SN of Spark100-Ef1a-Parkin, Spark100-CAG-Parkin, Spark100-hSyn1-Parkin, Spark100-TH (full)-Parkin, Spark100-TH (mini)-Parkin. Statistically significant pairwise comparisons between the controls (naïve or diluent) and the experimental groups were reported as follows: ****p<0.0001. Statistically significant pairwise comparisons between experimental groups were reported as follows: ^####^p<0.0001.

## Discussion

Our research provides a foundation for the development of AAV- Parkin gene therapy as a treatment for early-onset Parkinson’s disease (EOPD) caused by biallelic loss of function mutation in *PRKN*. A key advantage of gene therapy is its ability to target specific cells, which in this case allows for the direct delivery of functional Parkin to the dopaminergic cells of the substantia nigra. This precise delivery is crucial for restoring proper clearance of damaged mitochondria via mitophagy, a process critical for slowing the progression of dopaminergic neuron loss.

We have demonstrated that replenishing functional Parkin successfully restores pUb^Ser65^ defects in a range of in vitro models, including patient-derived cells. Furthermore, our in vivo studies establish a feasible and well-tolerated strategy for the targeted expression of Parkin specifically within SN- an approach gaining significant traction due to its demonstrated effectiveness in treating conditions like aromatic L-amino acid decarboxylase (AADC) deficiency^14^.

### Developed a model to study Parkin Activation and Mitophagy initiation In Vitro

A critical aspect of our study was our reinforcing pUb^Ser65^ as a reliable marker of Parkin activity. By inducing mitochondrial stress with FCCP, we demonstrated that Parkin KO neuroblastoma cells exhibit a deficiency in pUb^Ser65^ amplification, a deficit functionally rescued by the introduction of AAV-Parkin. The observed MOI-dependent increase in pUb^Ser65^, coupled with the reduction in Parkin protein levels upon stress, strongly supports the notion that AAV-delivered Parkin actively engages with and is degraded alongside dysfunctional mitochondria via mitophagy. Crucially, the direct visualization of AAV-Parkin co-localizing with mitochondria following FCCP treatment provides confirmation of its functional recruitment (Fig 1).

Extending these findings to EOPD patient-derived fibroblasts with diverse *PRKN* mutations reinforces the translational relevance of our work. The observation of diminished pUb^Ser65^ amplification in patient cells under valinomycin-induced mitochondrial stress, and its subsequent restoration with AAV-Parkin, demonstrates that our gene therapy approach can restore Parkin function in the setting of diverse variants (Fig 2).

### Demonstrated Transgene Expression and Tolerability in Mammalian Models

Demonstrating successful transgene expression and tolerability in a mammalian model is paramount for gene therapy development. We used AAV1 capsid in this study for its demonstrated tropism in brain tissues^20, 21^. Our rat study confirmed the feasibility of AAV-Parkin (human) delivery to the SN via intraparenchymal injection, achieving dose-dependent human Parkin expression that co-localized effectively with TH-positive dopaminergic neurons (Fig 3B). Quantitatively, the 3e8 vg/SN dose successfully achieved Parkin expression within our targeted range of 50-100% of endogenous levels, a crucial finding for therapeutic replacement strategies. At the 3e9 vg/SN and 3e10 vg/SN doses, we achieved Parkin overexpression significantly beyond physiological levels (data not shown). We observed some variability in Parkin expression (Fig 3C and 3D). This could be because we were targeting a small, deep brain structure like the SN, but it didn’t affect the study’s overall conclusion.

While doses below 3e10 vg/SN were well-tolerated, the highest dose resulted in significant upregulation of glial activation markers (Gfap and Iba1), as well as a reduction in striatal dopamine and impaired dopamine turnover, indicative of dopaminergic neuronal dysfunction (Fig 3E-H). Although a vector-only control at this highest dose was not included, the correlation between supraphysiological Parkin expression and adverse glial activation strongly suggests excessive Parkin levels, rather than just viral load, can induce a neuroinflammatory response and neuronal compromise. This underscores the delicate balance required for effective gene therapy and highlights the necessity of precise dose-finding to avoid unintended consequences, even with a protective factor like Parkin.

### Promoter Selection for Optimal Targeting

In parallel with the rat study, we conducted a complementary study in mice to select the optimal promoter for our proprietary Spark100 capsid, aiming for the best transduction of target cell populations. Spark100 showed similar transduction of dopaminergic neurons in the SN as the commonly used AAV1 and AAV9 capsids (data not shown). A dose of 1e10 vg/SN was selected for this study to account for the differential strengths of the promoters used (hSyn1 is known to drive weaker expression than the CAG promoter). The objective was to ensure sufficient human Parkin expression from all promoters at the given dose and time point, thereby facilitating an informed interpretation regarding the optimal promoter selection.

The CAG promoter drove Parkin to supraphysiological levels of overexpression, causing significant gliosis (Fig 5A-C). While we didn’t see a significant reduction in TH levels in the SN and striatum compared to control animals, there was a consistent trend. This difference was only statistically meaningful when we compared Spark100-CAG-Parkin (human) to Spark100-hSyn1-Parkin (human) in the striatum (Fig 4E). Unlike the rat study (Fig 3), the mouse study had additional groups injected with vectors using Spark100 capsid at 1e10vg/SN (Fig 4B). Since these groups showed no signs of gliosis (based on Iba1 and GFAP analysis) compared to diluent and naïve groups, we can conclude that the significant gliosis seen in the Spark100-CAG-Parkin group was caused by the supraphysiological levels of human Parkin, and not by titers/dose of the capsid.

Other tested promoters were determined to be less optimal for more nuanced reasons. For instance, the TH promoter initially appeared to be an ideal candidate for expression in TH-positive neurons. However, we observed a loss of TH protein expression in dopaminergic neurons by immunohistochemistry (Fig 4C), without any associated upregulation of inflammatory markers. Ultimately, the hSyn1 and Ef1a promoters demonstrated the most ideal expression level and specificity for Parkin gene therapy, without any glial inflammation.

These findings collectively lay a foundation for a viable therapeutic approach for Parkin-PD, while also underscoring the need for dose-finding studies to optimize Parkin expression levels for clinical translation.

## Conflict of interest

SB, TGD, NG, AK, BW, EK, DMC, PGA, MN, RG, MGB, and E.R. are either present or past paid employees of Spark Therapeutics, Inc., a Roche company, and may hold equity in the organization.

This research was supported in part by the Intramural Research Program (ZIA NS003169) of the National Institutes of Health (NIH), National Institute of Neurological Disorders and Stroke (NINDS).

## Acknowledgments

We thank Xin Zhang, Lihua Shi, Bin Zhong and Charmaine Azutillo for vector support; FD Neurotech and Bo Wang for histology assistance; Aidan Smith, and Karthik Ramanathan for their contribution in the initial mouse studies. We would also like to extend our gratitude to all the patients for donating the skin fibroblasts used in this study. The graphical abstract was generated using Biorender.

## Financial Disclosures

The contributions of the NIH author(s), DPN and AJG were made as part of their official duties as NIH federal employees, are in compliance with agency policy requirements, and are considered Works of the United States Government. However, the findings and conclusions presented in this paper are those of the author(s) and do not necessarily reflect the views of the NIH or the U.S. Department of Health and Human Services.

## Authors’ Contribution

SB, TGD, DPN, MGB and LR conceptualized the project, designed the studies and experiments. DPN, PGA, RG, MGB, and ER supervised the research. SB, TGD and DMC designed the plasmids. SB, TGD, NG, AK, AJG, conducted *in vitro* experiments and analyzed data. MN, RG, BW, EK, NG, TGD, developed methodologies and conducted the investigation of animal samples. TGD, NG, SB, DPN, MGB, and E.R. contributed to writing and editing. All authors reviewed and edited the manuscript.

## Supplemental data file

**Supplementary Figure 1:**
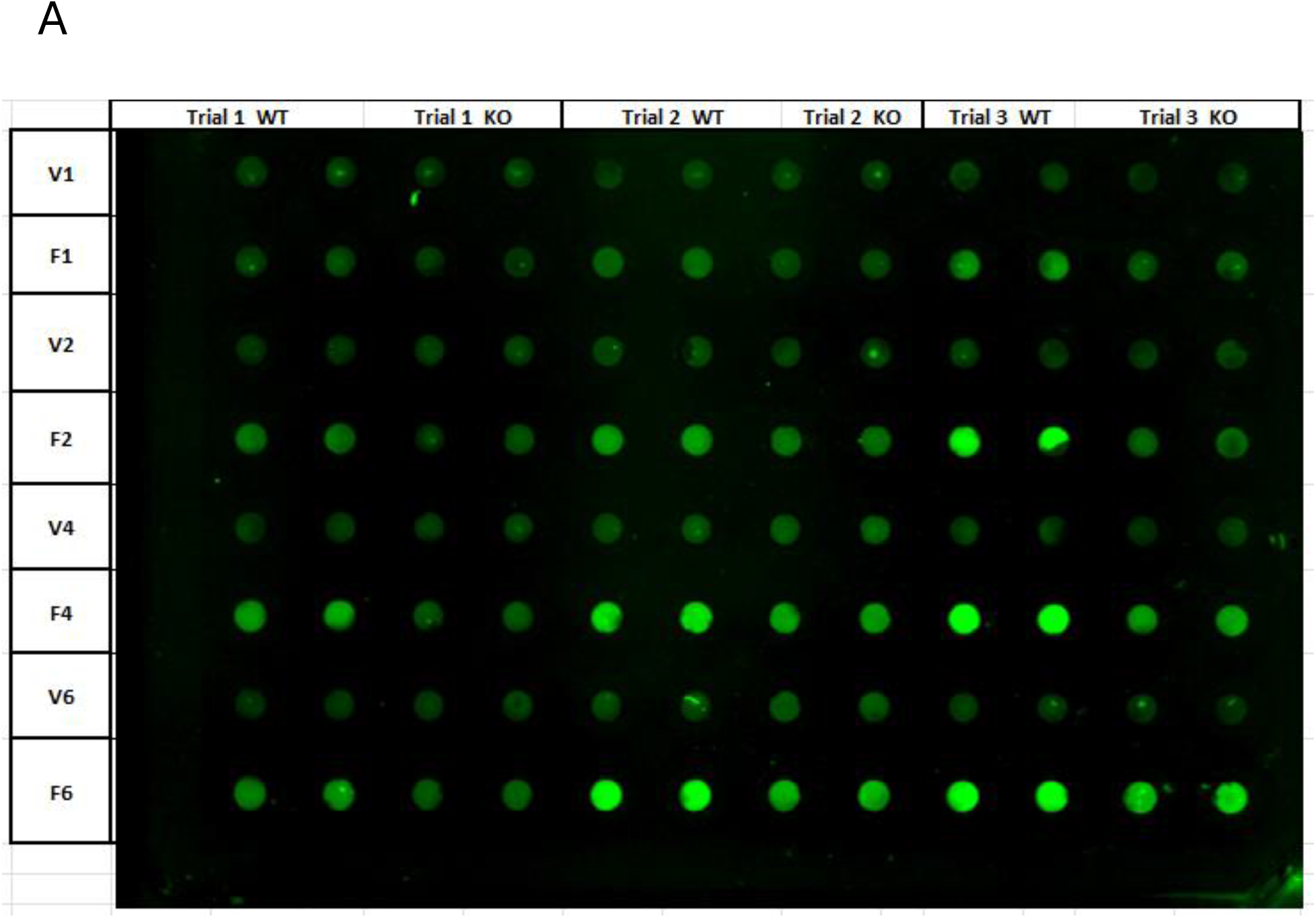
Identification of Phospho-ubiquitin serine 65 (pUb^Ser65^) as a marker of Parkin-mediated mitophagy: A) Dot blot image measuring pUb^Ser65^ response in WT and Parkin KO SH-SY5Y cell lines. Across 3 trials, FCCP treatment triggered a significant upregulation in pUb^Ser65^ response across a 6-hour time course in WT.

**Supplementary Figure 2:**
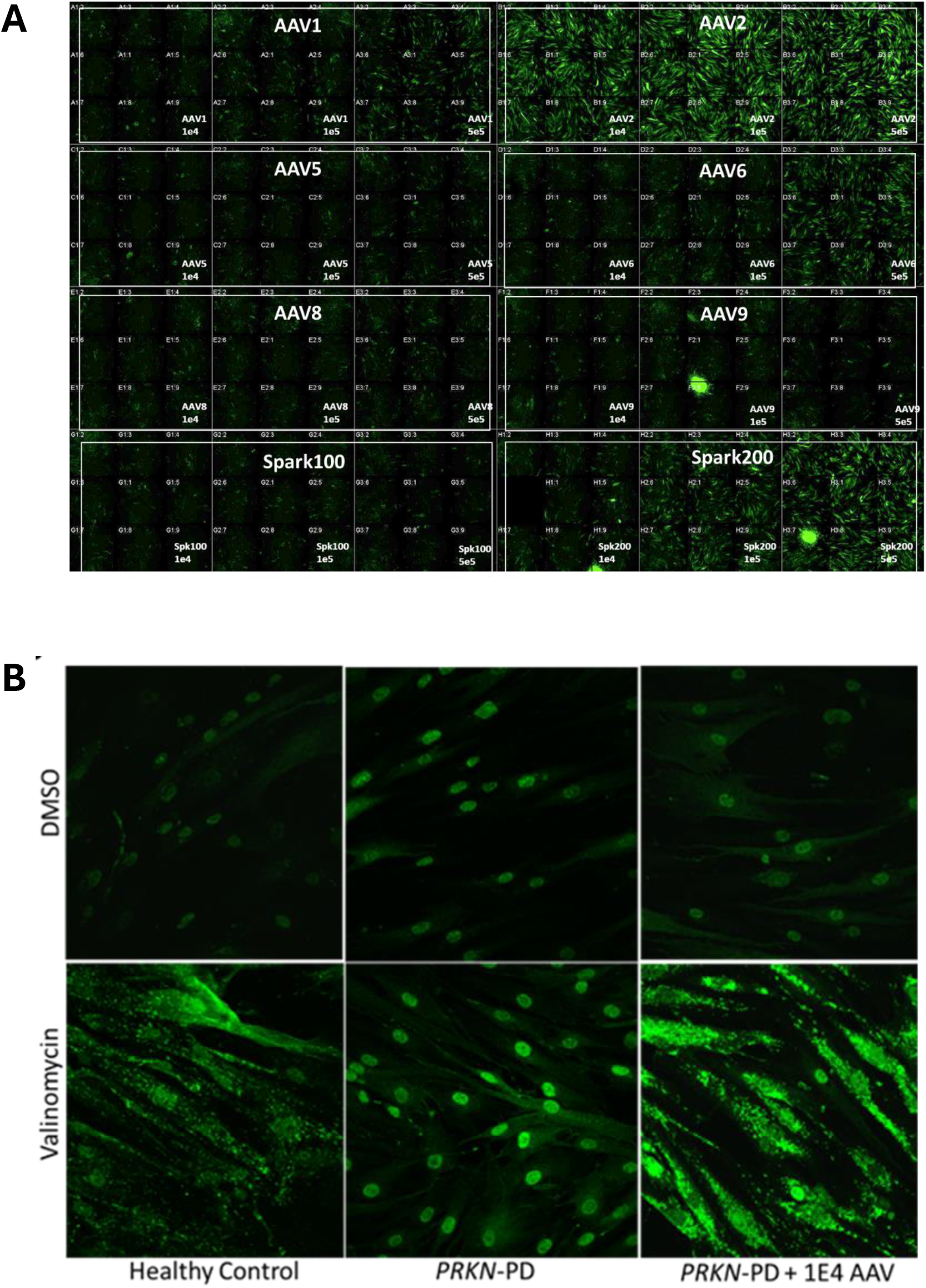
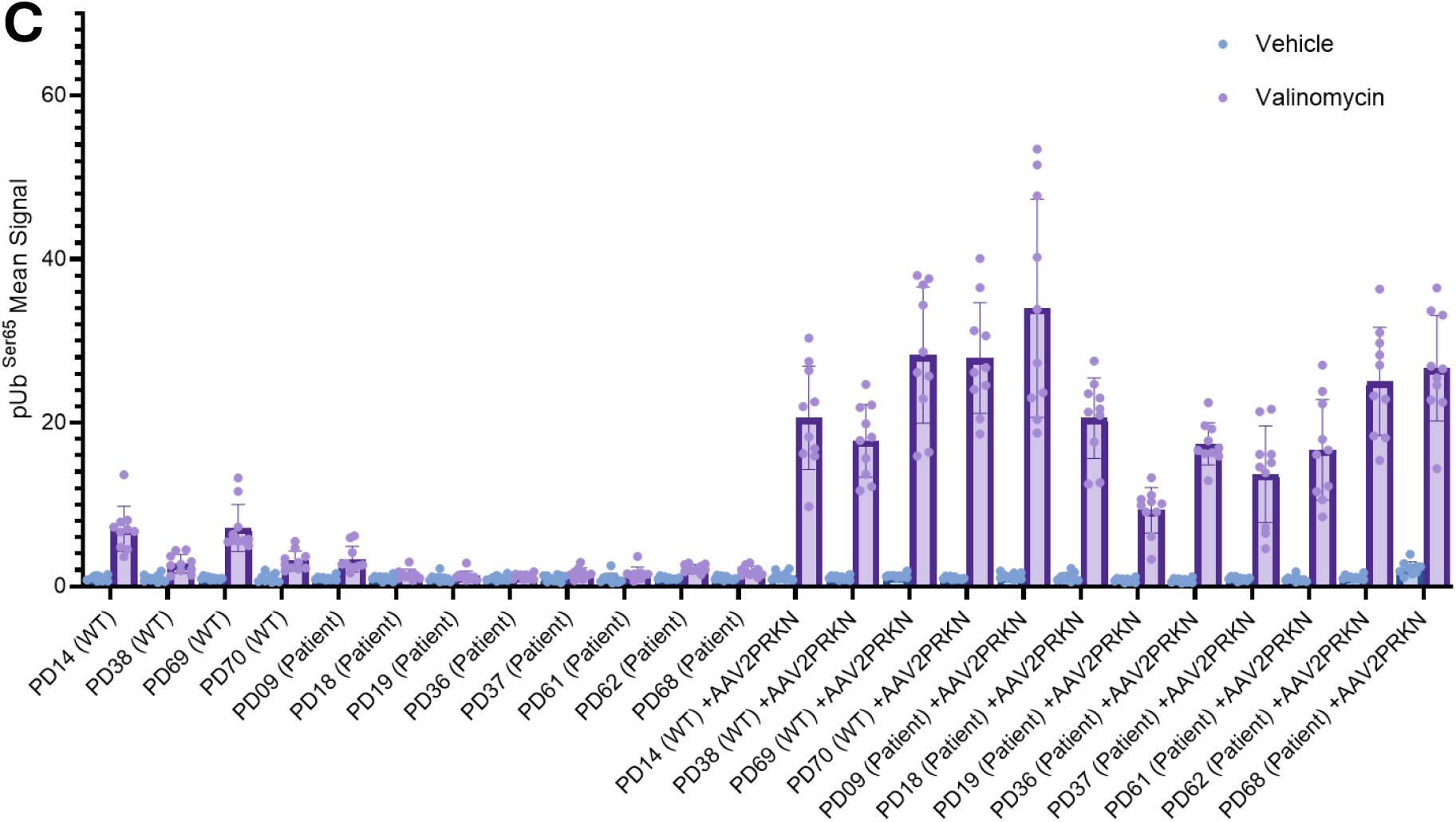
AAV2-CAG-Parkin (human) successfully transduces and restores pUb^Ser65^ signal in PD patient fibroblasts: A) Representative images of WT human skin fibroblasts tested with various AAV capsids expressing GFP. B) Representative images of pUb^Ser65^ staining in PD patient-derived fibroblasts showing restoration of mitophagy when cells are treated with AAV2-CAG-Parkin and stressed with valinomycin c) Quantification of pUb^Ser65^ fluorescence mean signal intensity from each Healthy control and PD patient lines, with and without valinomycin treatment.

**Supplementary Figure 3.**
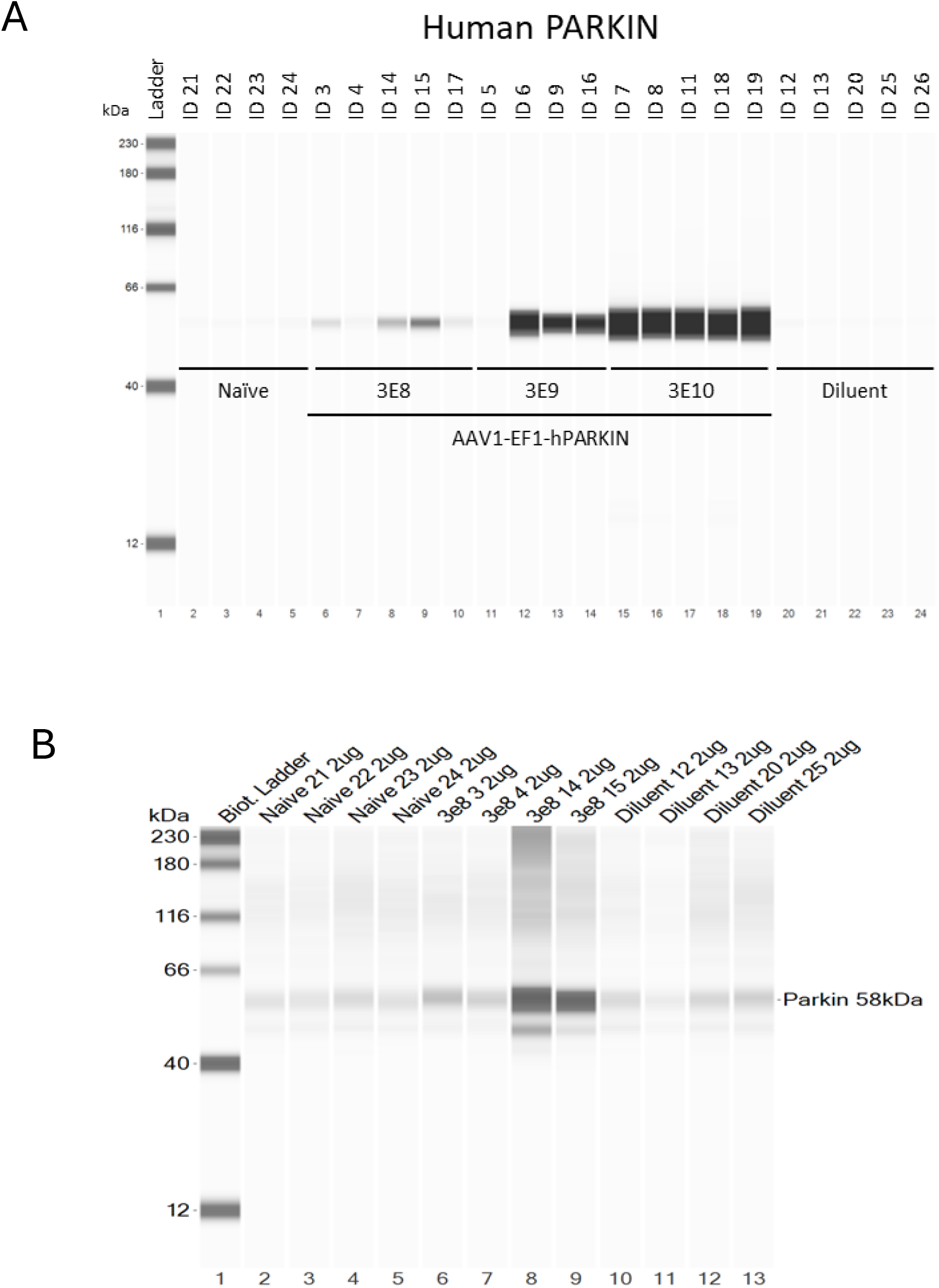
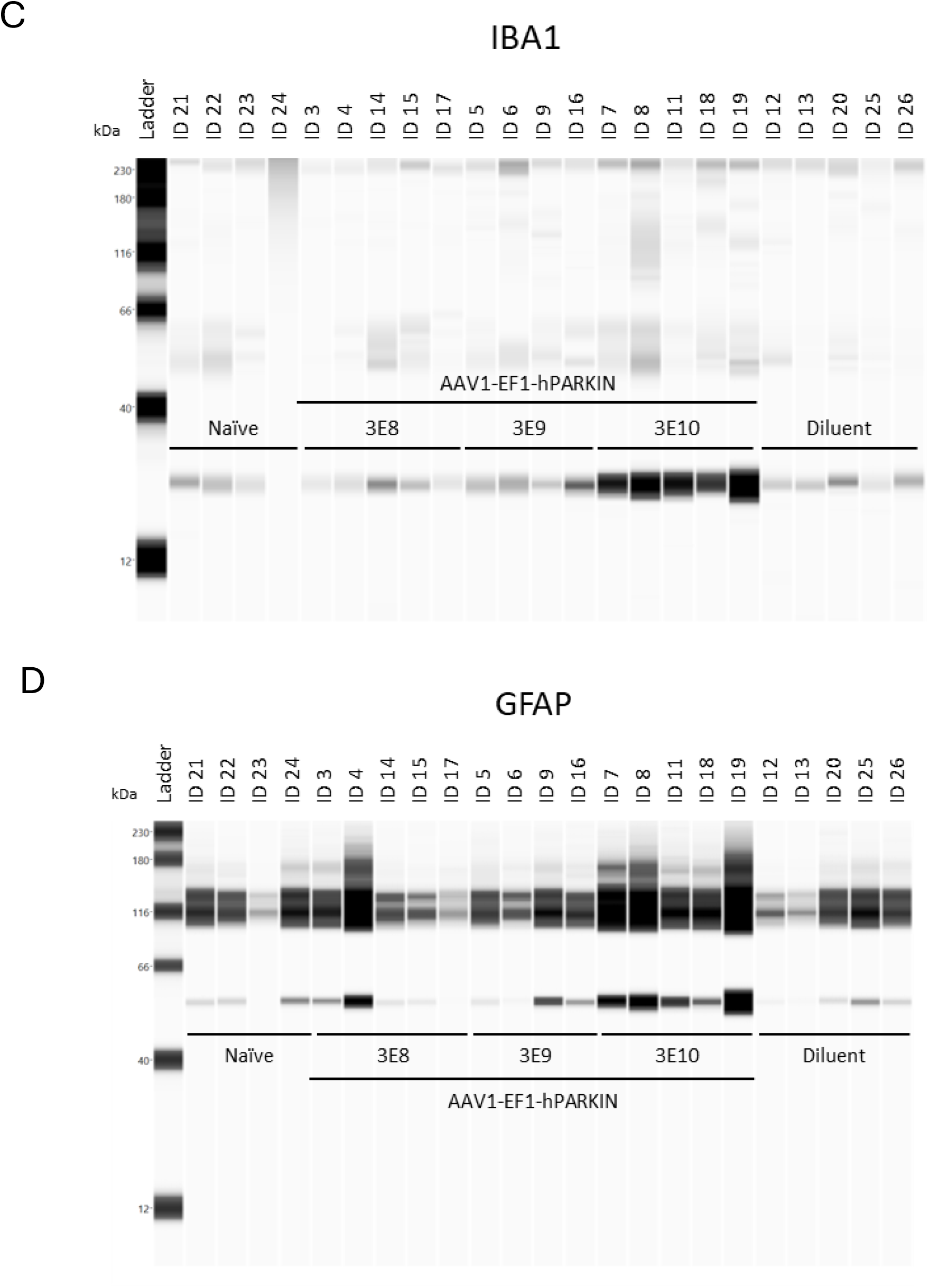
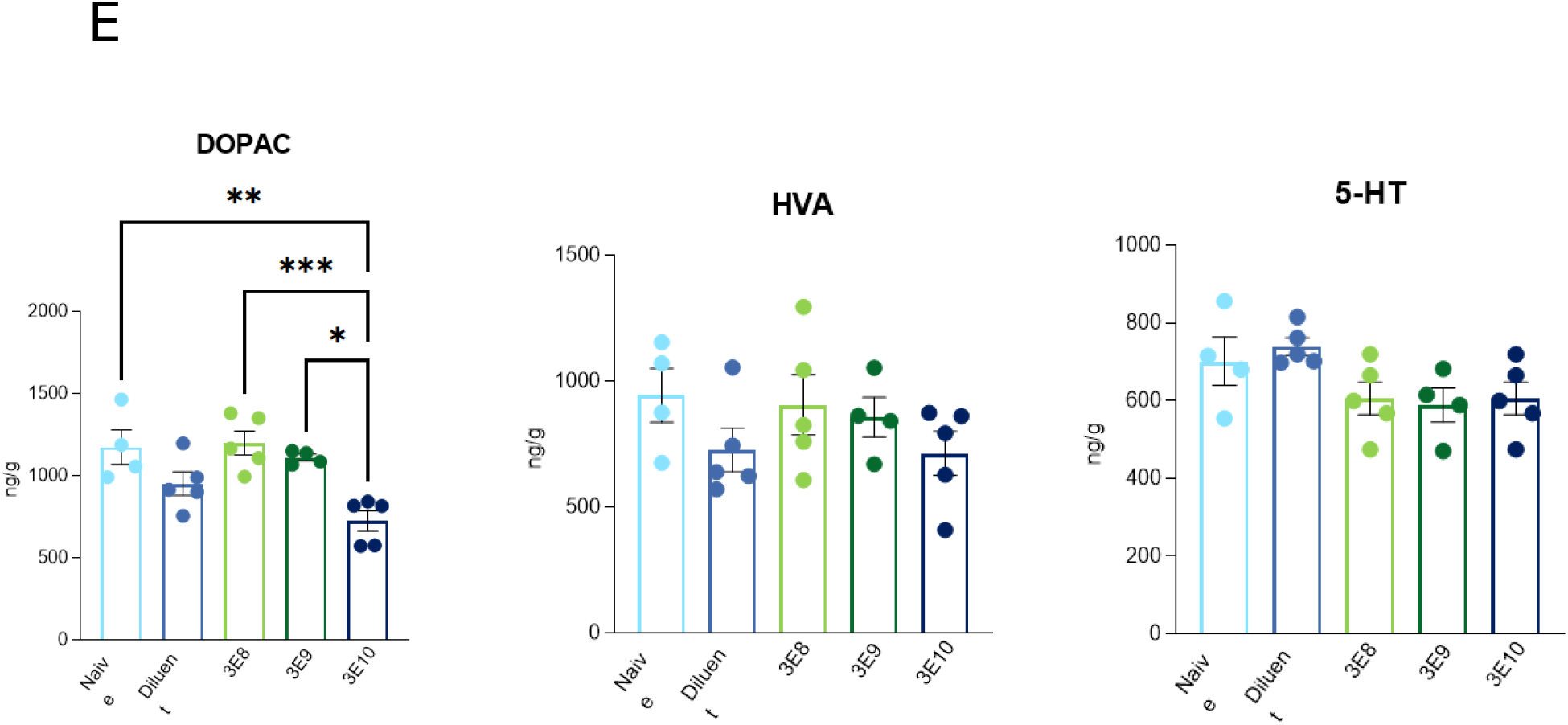
Dose-dependent Parkin (human) expression following stereotaxic delivery of AAV-Ef1a-Parkin (human) in Rat substantia nigra pars compacta (SNpc): Capillary electrophoresis images showing expression of human Parkin (A), Total Parkin (B) GFAP (C) and Iba1(D) from SN tissues of individual animals following stereotaxic delivery of AAV-Ef1a-Parkin (human) in Rat substantia nigra pars compacta (SNpc) (E,F) HPLC for dopamine metabolites 3,4-dihydroxyphenylacetic acid (DOPAC), homovanillic acid (HVA) and 5-HT.

**Supplementary Figure 4:**
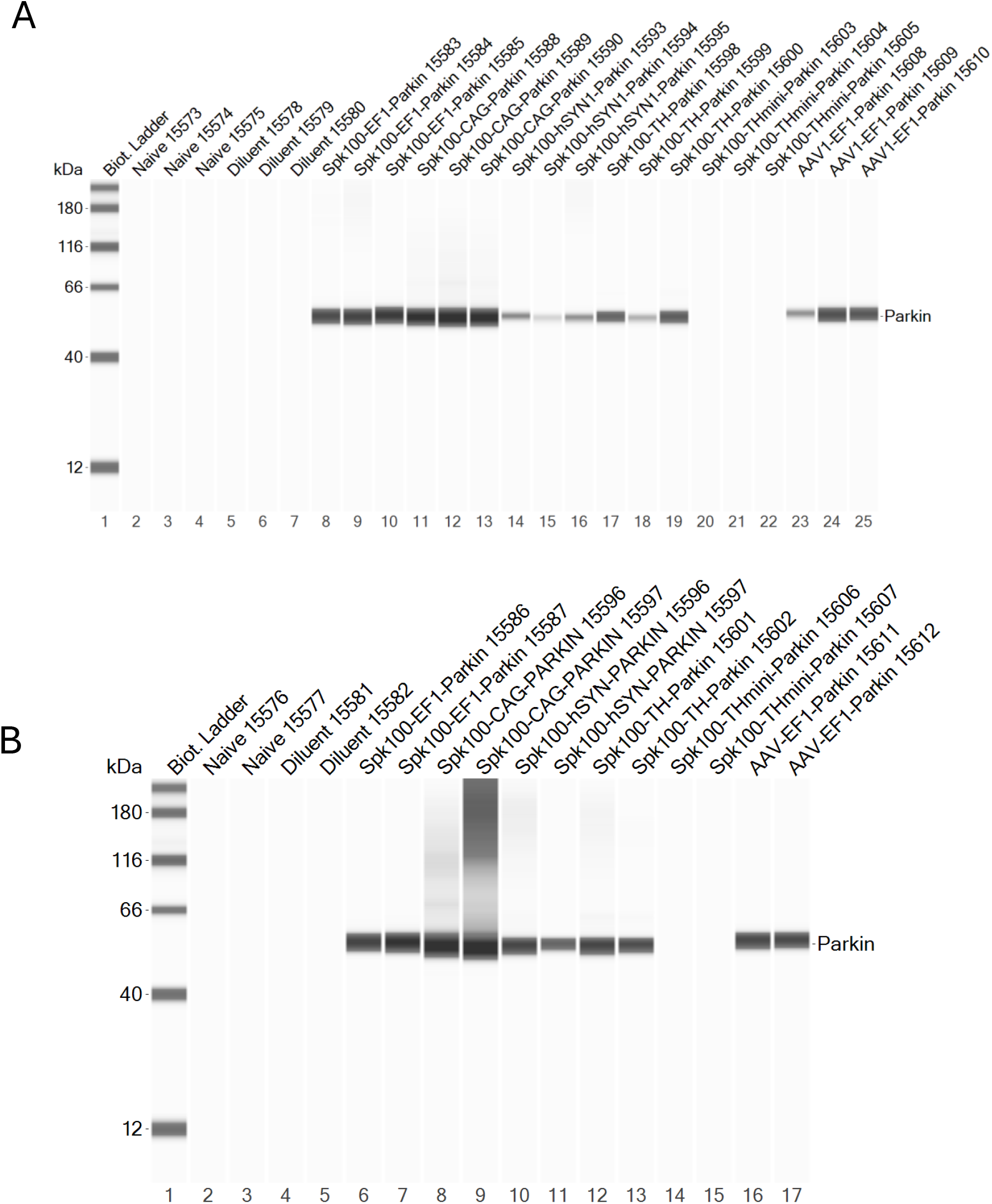
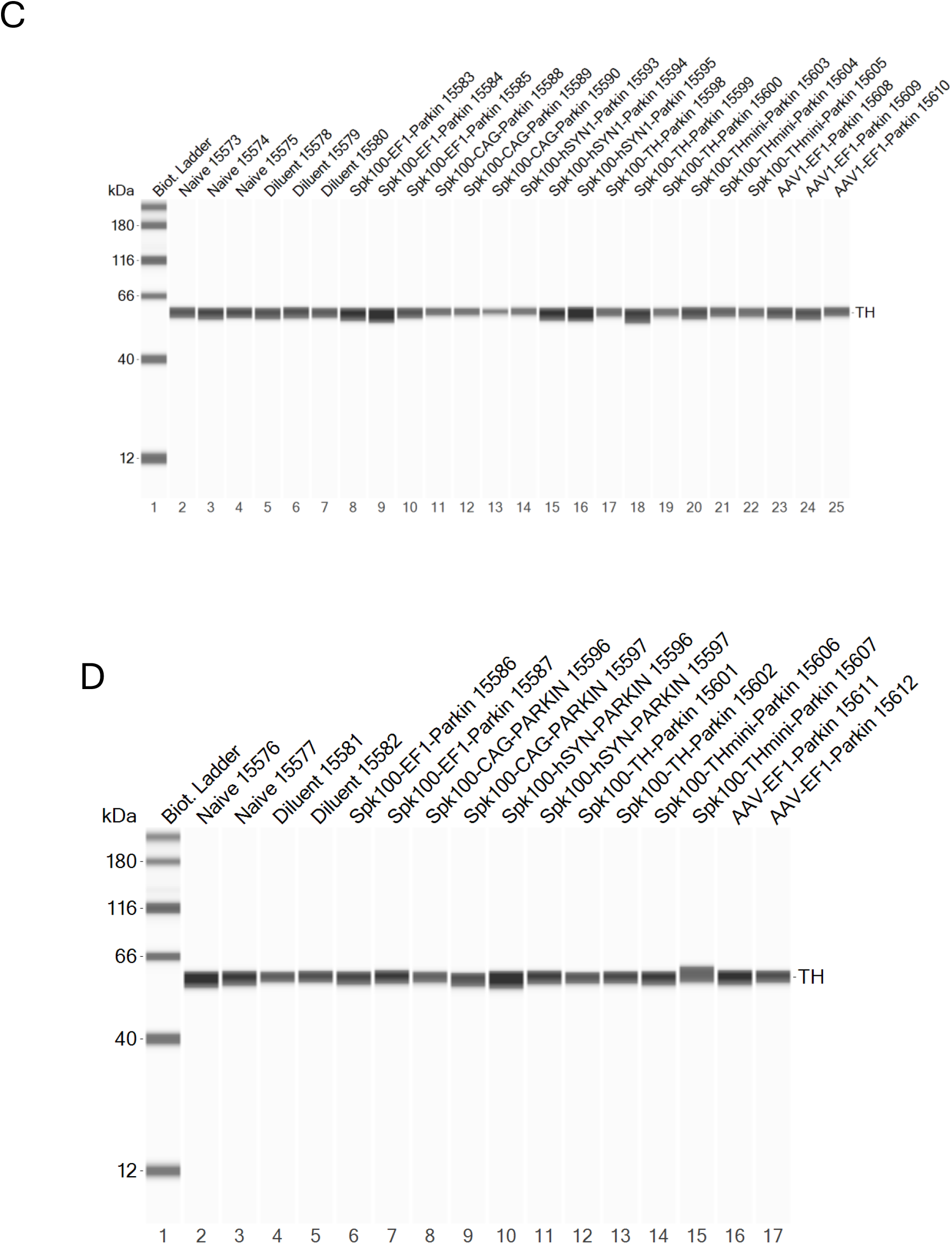
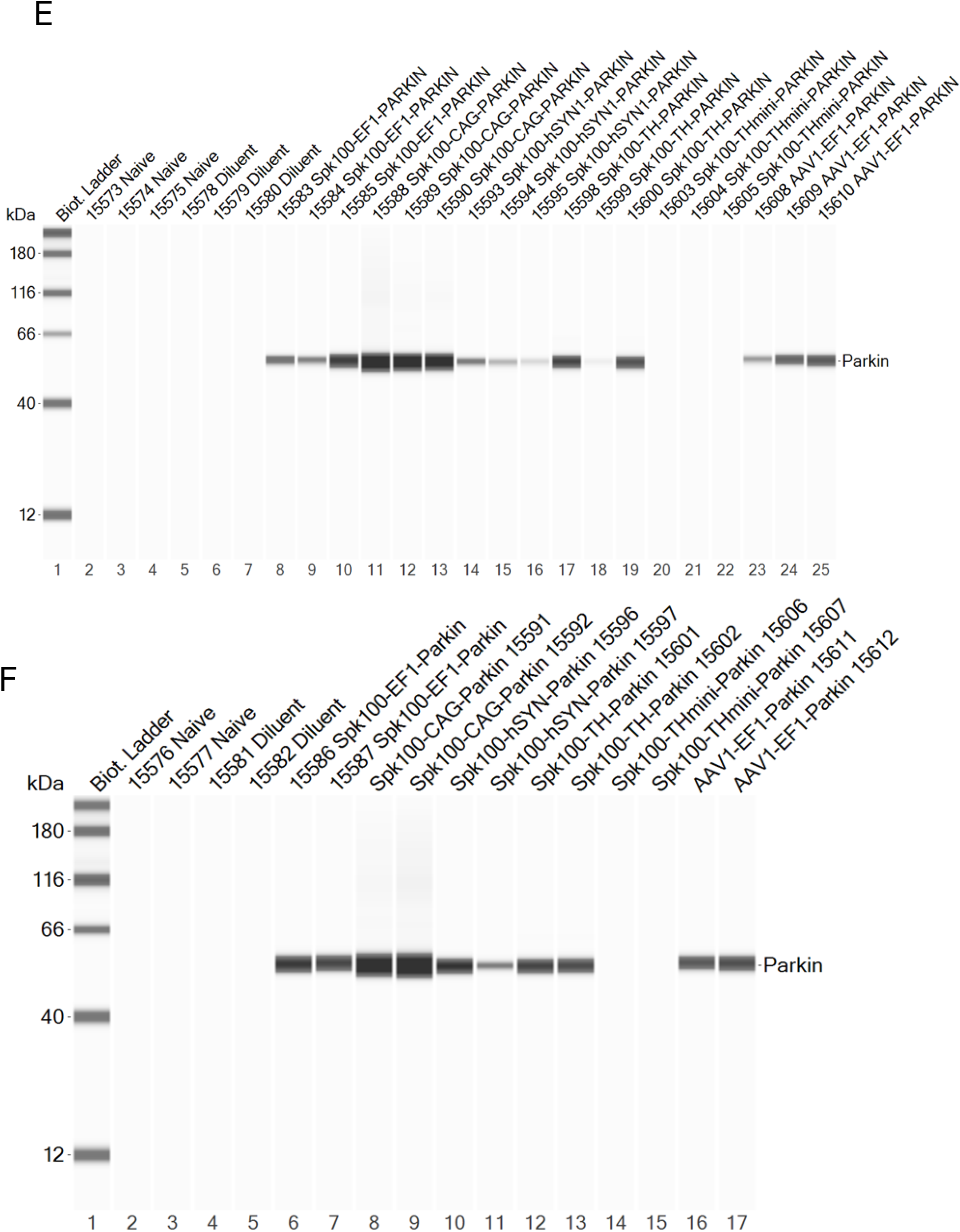
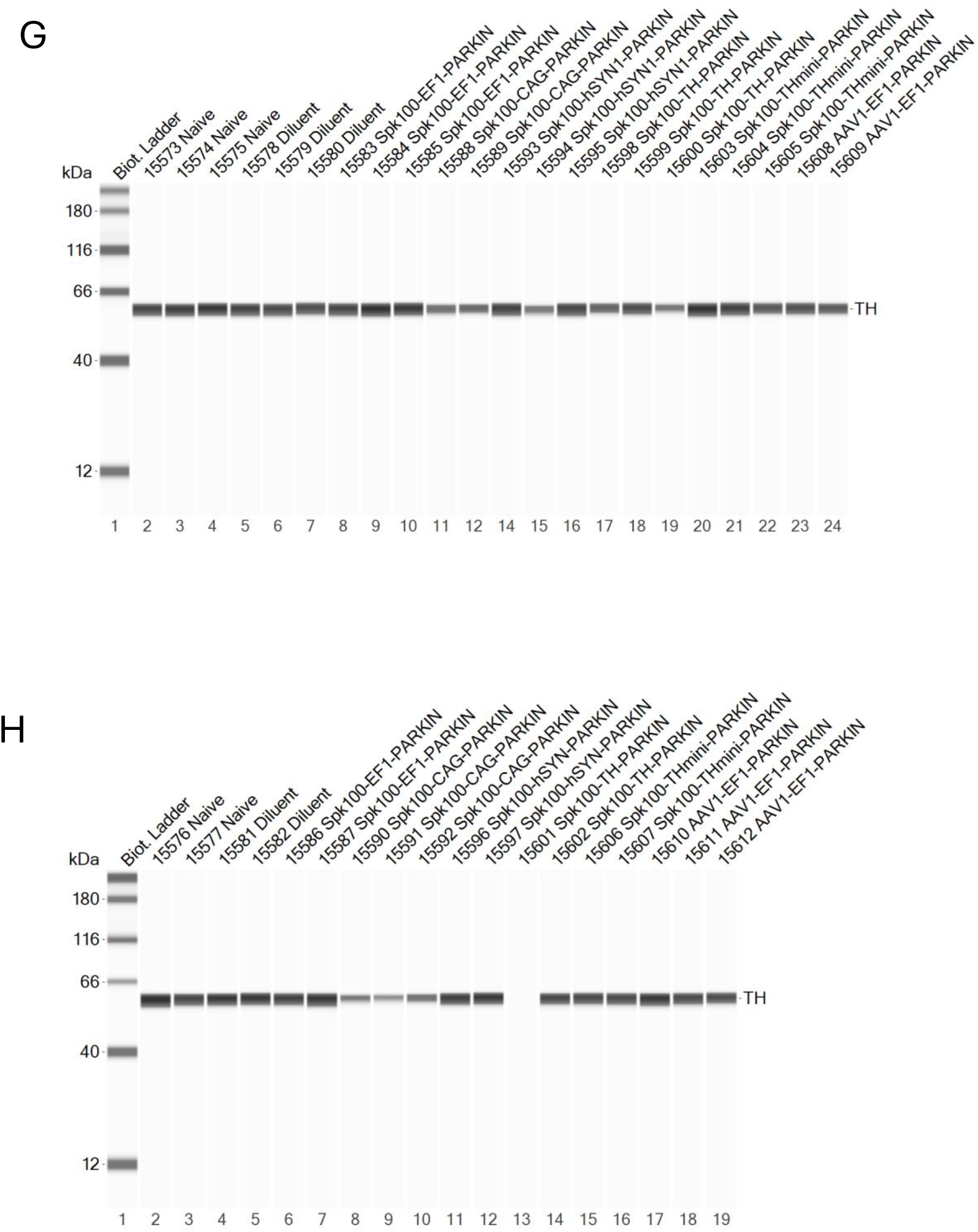
Optimal Promoter selection for expressing Parkin(human) Expression in dopaminergic neurons of mouse SN. Capillary electrophoresis images showing human Parkin and TH expression of individual animals from SN (A-D) and striatum (E-H) following stereotaxic delivery of 1e10 vg/SN of Spark100-Ef1a-Parkin, Spark100-CAG-Parkin, Spark100-hSyn1-Parkin, Spark100-TH (full)-Parkin, or Spark100-TH (mini)-Parkin to the SN.

**Supplementary Figure 5:**
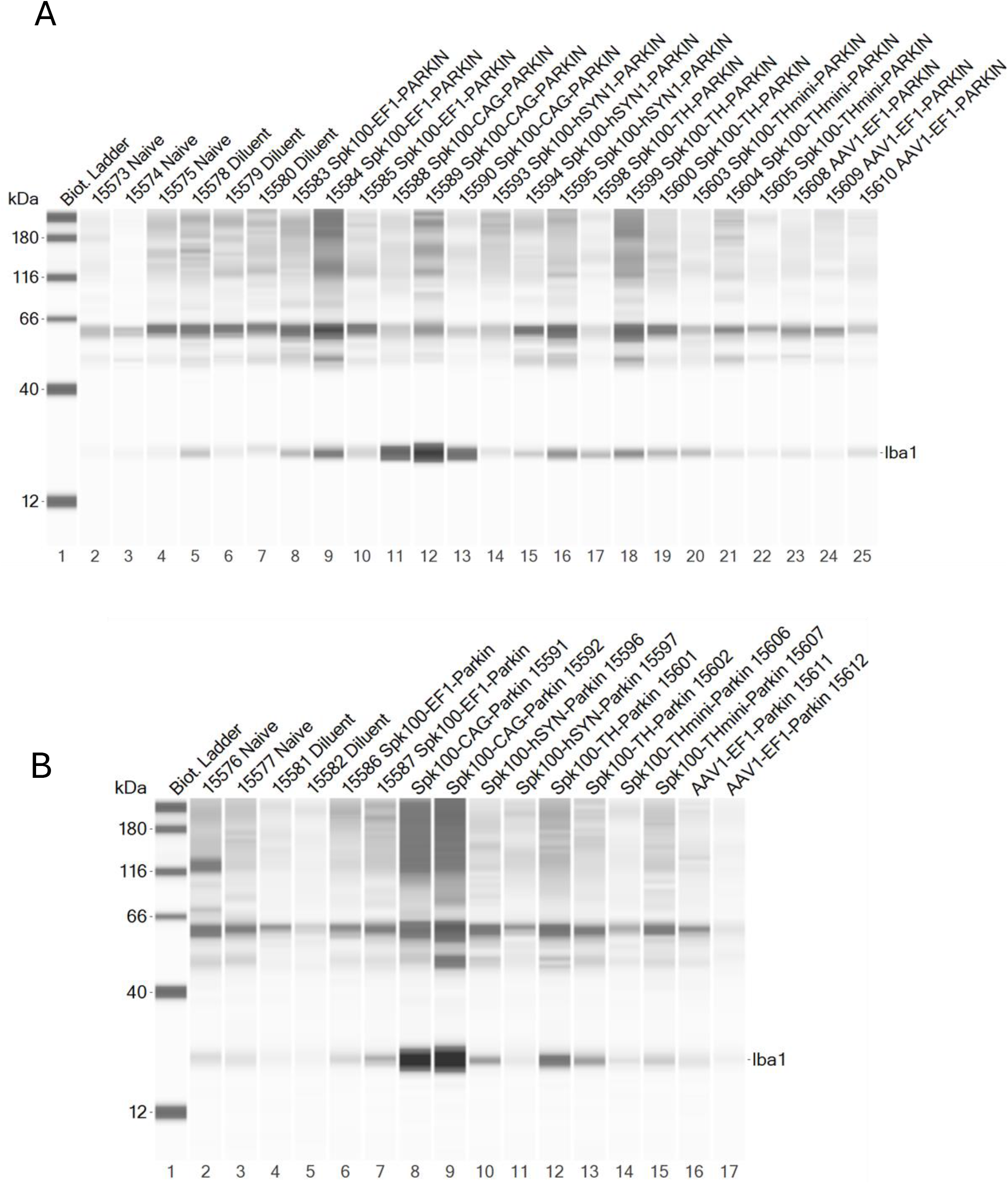
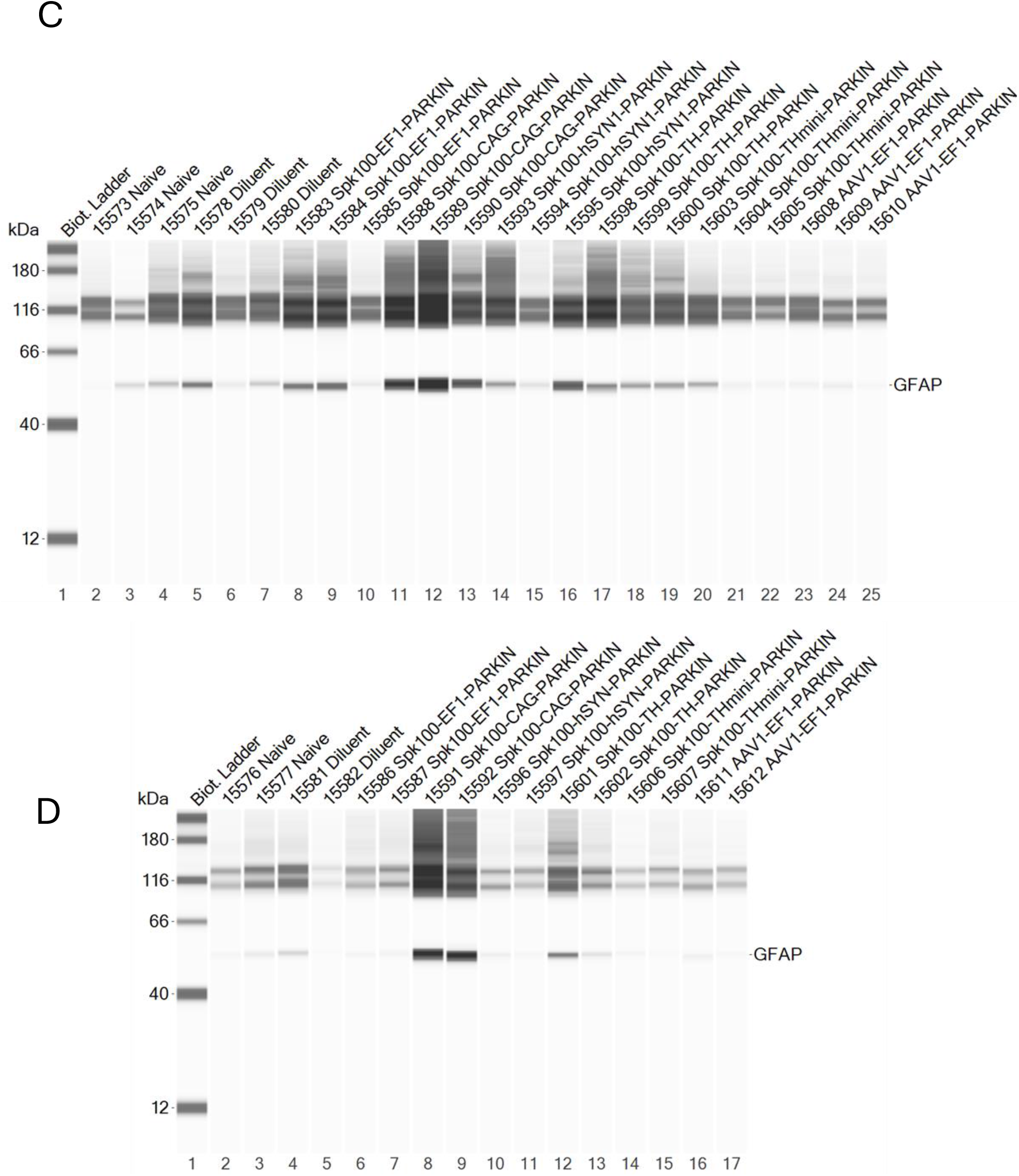
Evaluation of gliosis markers Iba1 and GFAP in mouse SN Tissues following overexpression of Parkin (human) from different promoters. Capillary electrophoresis images showing Iba1 (A,B) and GFAP (C,D) expression of individual animals from SN following stereotaxic delivery of 1.0×1010 vg/SN of Spark100-Ef1a-Parkin, Spark100-CAG-Parkin, Spark100-hSyn1-Parkin, Spark100-TH (full)-Parkin, or Spark100-TH (mini)-Parkin to the SN.

## Supplementary Materials and Methods

### In-vitro studies

#### pUb^Ser65^ dot blot from SH-SY5Y cells

Wild-type and Human Parkin knock out SH-SY5Y neuroblastoma cells were obtained from abcam (cat #ab280042). The control wild-type SH-SY5Y cells were provided with the Human Parkin knock out cells (cat #ab275475). SH-SY5Y cells were cultured and maintained as per manufacturer’s instruction, with the exception of DMEM F-12 media (Gibco, 11320033) being used instead of EMEM.

WT and Parkin knock out SH-SY5Y cells were plated in 6 well plates at 1e6 cells per well and treated with 20uM FCCP (Cayman Chemical, Item# 15218) diluted in Dimethyl sulfoxide (DMSO; Sigma-Aldrich, Catalog #D2650) or vehicle (DMSO) for 1, 2, 4 and 6 hours. Post-treatment the cells were collected, and the extracted protein was quantified using standard BCA assay (Thermo Fisher, Cat. No. 23225). 2.5ug of protein was diluted in a total of 25ul using tris-buffered saline (TBS) and used for a dot-blot analysis of pUb^Ser65^. The antibody details are included in the table below.

In brief, 1ug diluted protein (10uL) lysate was loaded onto a nitrocellulose membrane on a microfiltration blotting device with an attached vacuum manifold base (BioRad Cat #1706545). Samples were incubated on the membrane at RT for 1 hour. Blocking solution (LiCor Intercept Cat#927-80001) was added and incubated on an orbital shaker for 1 hour at RT. Primary antibody solution (1:500 Phospho-Ubiquitin Ser65 Cell Signaling Cat# 62802 diluted in 20mL 0.1% Tween blocking solution) was added and incubated overnight with gentle rocking at 4C. The following day, 3 washes with TBST (TBS+0.1% Tween) were performed. Secondary antibody solution (Licor IR800 Cat# 926-32213 diluted 1:20,000 in blocking solution) was added and incubated with gentle rocking for 1 hour at RT. Membrane was scanned on LiCor Odyssey for blot quantitation.

#### SH-SY5Y localization of Parkin and mito-RFP upon FCCP treatment

SH-SY5Y cells (WT and Parkin knock out) were plated on ready-to-use PDL coated 96-well plates (Perkin Elmer) at 20,000 cells/well on Day 0. Depending on the treatment paradigm on day 1 cells were either treated with Diluent or AAV1-Ef1a-Parkin (human) (stock titer 1.04e13vgs/ml) at multiplicity of infection (MOI) of 1e5vp (vector particles)/cell for 20,000 cells. On Day 2, mitochondria were tagged using mito-RFP (catalog #C10601 Invitrogen) at a MOI of 40 particles/cell. 24 hours after the labeling of mitochondria, cells were treated with 20uM of FCCP (stock concentration of 10mM) for 4hrs. After treatment with FCCP, cells were washed and fixed with 4% PFA for 15mins. This was followed by a standard immunocytochemistry protocol. Briefly, after the fixation step, cells were washed and permeabilized with PBS+0.5%TritonX100 for 15mins, washed, and blocked for 2hrs at RT with blocking buffer (PBS+0.1%Tween 20+10%Goat serum). Post blocking and washing, cells were incubated with Primary rabbit anti pUb^Ser65^ (Cell-signaling Cat #62802) at 1:250 or rabbit anti Parkin (Invitrogen Cat #702785) at 1:500 dilution overnight in Antibody buffer (PPBS+0.1%Tween 20+1%Goat serum) at 4C. The next day, primary antibody was aspirated and cells were washed with a washing buffer (PBS+0.1%Tween 20) 5 times, 5 minutes each wash and incubated with Goat anti Rabbit conjugated to Alexa Fluor 488 (GFP) for 1hr at RT. Cells were washed, treated with DAPI (1:1000) and imaged using Opera Phenix. Images were acquired using 10X air objective for EGFP, Cy3 and DAPI channel. The entire well image was stitched together to qualitatively assess parkin (EGFP) colocalization to mitochondria (Cy3), resulting yellow pixels post FCCP treatment.

#### Patient derived fibroblast ICC for pUb^Ser65^

Healthy control and Parkin-mutation (PRKN-PD) fibroblasts were obtained from NINDS. All participants gave written informed consent according to the Declaration of Helsinki to protocols approved by the Institutional Review Board of NINDS before undergoing research procedures. A detailed list of the specific mutations in the PRKN gene are listed in the results section. Patient fibroblast lines were established from 3-mm punch biopsies taken from the forearm. Cell lines were assayed at passage 11 or before. Cells were cultured in high glucose DMEM (Life Technologies, cat#11995073) with 10% fetal bovine serum.

For selection of optimal transduction conditions in fibroblasts, wild-type fibroblasts were seeded at 20000 cells/well in a 96-well plate format. Next day, the cells were transduced with 3 MOI of AAV-vectors (1e4, 1e5 or 1e6 vp/cell) expressing GFP from a CAG promoter (titers of each vector provided in the table below). 72hours post-transduction, the efficiency was assessed by imaging using Opera Phenix.

12 lines of primary patient-derived fibroblasts were seeded in Ibidi slides (cat# 80827-90) at 20,000 cells per well. The next day cells were transduced with AAV2-CAG-Parkin (human) at an MOI of 1e4 vp/cell. After 72 hours of transduction, cells were treated with 1uM Valinomycin (Valinomycin, solid (SIGMA Cat#V0627-25MG) or an equivalent volume of vehicle (DMSO). After 3 hours of Valinomycin treatment fibroblasts were fixed and stained for pUb^Ser65^following standard immunocytochemistry protocol. The antibodies used for this assessment are described in the table below. Slides were imaged using an Olympus FV3000 and a 40X silicon oil immersion objective. Images were analyzed using ImageJ. For analysis, individual cells were outlined. An ROI of the mitochondrial region was created by converting the Cytochrome *c* slice into a binary mask. Measurements of the pUb^Ser65^ slice were taken within this mask. The mean grey value of pUb^Ser65^ was reported from individual cells.

**Table 1:** Table showing a list of Healthy controls and PRKN-PD patient fibroblast lines, with specific Parkin mutations (s), disease status and age of onset of disease.

| Sample Name | PD | Mutation | PD AAO | Sex |
| --- | --- | --- | --- | --- |
| FPD014 | No | Does not have biallelic mutations in Parkin | N/A | M |
| FPD038 | No | Does not have biallelic mutations in Parkin | N/A | M |
| FPD069 | No | Does not have biallelic mutations in Parkin | N/A |  |
| FPD070 | No | Does not have biallelic mutations in Parkin | N/A | M |
| FPD009 | Yes | Parkin homo c.1244C>A; p.Thr415Asn | 27 | F |
| FPD018 | Yes | Parkin compound het (Exon11 del; p.G429D) | 31 | M |
| FPD019 | Yes | Parkin compound het (Exon11 del; p.G429D) | 37 | F |
| FPD036 | Yes | Parkin compound het (c.167 T>A, p.Val56Glu;<br>Partial gene deletion including E7 | 35 | F |
| FPD037 | Yes | Parkin compound het (c.167 T>A, p.Val56Glu;<br>Partial gene deletion including E7 | 32 | F |
| FPD061 | Yes | Parkin compound het (p.R275W, del E3-4) | 19 | M |
| FPD062 | Yes | Parkin compound het (p.R275W, del E3-4) | 37 | M |
| FPD068 | Yes | Parkin compound het VUS splice site<br>(rs1204087943) and E3-4 deletion | 27 | F |
| FPD075 | Yes | Parkin compound het E3 del and E3-E4 del | 37 | M |

#### Vector Production

Recombinant AAV vectors were produced at the Research Vector Core at Spark Therapeutics Inc. by transfecting adherent HEK293 cells using a calcium-phosphate-based triple plasmid system containing pHelper, pRepCap, and a transgene plasmid (for Parkin) at a 1:1:1 mass ratio. After 96 hours, the cells were harvested and lysed by sonication to release the viral particles. The lysate was then treated with benzonase to digest unwanted host cell DNA and RNA, and the AAVs were concentrated using polyethylene glycol (PEG) precipitation. For final purification, the vectors were isolated via CsCl density-gradient ultracentrifugation and then buffer-exchanged into a PBS/Pluronic F-68 solution through dialysis. The quality of the final product was thoroughly checked: vector genomes were quantified using TaqMan qPCR with BGH polyA-specific primers (forward 5′-CTTGCCTTCCTTGACCCT-3′, reverse 5′-CCCAGAATAGAATGACACCTACT-3′) and a probe (56-FAM/TTAGGAAAG/ZEN/GACAGTGGGAGTGGC/3IABkFQ); capsid protein purity was confirmed by SDS-PAGE with a Sypro Ruby stain; and endotoxin levels were verified to be less than 1 EU/mL. The final, quality-controlled vectors were stored at −80°C until they were needed ^16^.

#### Tolerability/Dose Ranging Study

The study included 24 male CD® Sprague Dawley rats (n=4/5 per group) aged 8 to 10 weeks at dosing (Day 0). Animals remained untreated (Group 1, n=4), received the diluent (Group 2, n=5) or the AAV1-Ef1a-Parkin (human), stock titer 1.04e13vgs/ml at dose 3.0×10^10^ (Group 5, n=5), 3.0×10^9^ (Group 4), or 3.0×10^8^ vg/SN (Group 3, n=5), and were monitored until necropsy 8 weeks post-dosing. The animals were randomized during surgical procedures to avoid any surgical bias. The rat tolerability study was conducted at Charles River Laboratories, Finland and was performed as specified in the license authorized by the national Animal Experiment Board of Finland and according to the National Institutes of Health (Bethesda, MD, USA) guidelines for the care and use of laboratory animals.

Twenty-four animals received bilateral intraparenchymal injections into the SNpc at the following coordinates: AP: 5.2 (posterior from bregma), ML: +/− 2.2, and DV: −7.5 (from the brain surface). AAV1-Ef1a-Parkin (human) were injected at a volume of 3uL/hemisphere using 10uL Hamilton syringe with a 26 G beveled needle. Animals were maintained under 2.5% isoflurane anesthesia for the length of the surgery. After surgery the skin was closed and disinfected. Animals were carefully monitored by lab personnel twice a day following surgery.

Rats were then transcardially perfused with ice cold heparinized (Heparin 2.5 IU/ml) saline to remove blood from the brains. The brains were removed carefully and were freshly dissected to be used for biochemical analysis (flash frozen, described in the section sample processing) or Immunohistochemistry (Fixed, described in the section immunostaining).

#### Promoter assessment

In-house mouse studies were approved by the Institutional Animal Care and Use Committee (IACUC) of Rowan University, Protocol No. 2020-1215 [2018-007]. Forty male animals (Mus musculus / C57BL/6) from Jackson Laboratories (Bar Harbor, ME, stock number# 000664) aged between 8-12 weeks received bilateral intraparenchymal (IPa) injections into the SNpc at the following coordinates: AP −3.1, ML ± 1.2, and DV −4.3, from the top of the skull and relative to the Bregma. The test articles (listed below in the AAV-vectors section) were injected at a volume of 2 µL per brain hemisphere and at a 0.4 µL/min rate. Animals were maintained under gas anesthesia using 3% isoflurane (via nose cone, at 1 L/min) throughout the procedure. Neosporin was applied after the stitches were placed to close the wound, and mice received carprofen (5 mg/kg subcutaneously). Animals were monitored by the Rowan University animal care staff on Days 1 and 2 post-surgery. Additional carprofen was given if needed.

On study Day 0, animals remained untreated (Group 1), received the diluent (Groups 2), or the following test articles at 1.0×10^10^ vg/SN: Spark100-Ef1a-Parkin (human) (Group 3), Spark100-CAG-Parkin (human) (Group 4), Spark100-hSyn1-Parkin (human) (Groups 5), Spark100-Th(full)-Parkin (human) (Groups 6), Spark100-TH(mini)-Parkin (human) (Groups 7) and AAV1-Ef1a-Parkin (human) (Groups 8). All animals received bilateral intraparenchymal (IP) injections into the SNpc and were monitored for up to 2 months (56 days) after injection.

On study Day 56, animals were sacrificed by live decapitation using a guillotine. The brains were removed carefully and were freshly dissected to be used for biochemical analysis (flash frozen, described in the section sample processing) or Immunohistochemistry (Fixed, described in the section immunostaining).

The sequence of the promoters is listed in the Supplementary files.

**Table 2:** List of vectors used for in-vitro Studies.

| Vector name | Titer (vg/ml) |
| --- | --- |
| AAV1-CAG-GFP | $4.72 \times 10^{12}$ |
| AAV2-CAG-GFP | $1.19 \times 10^{13}$ |
| AAV2retro-CAG-GFP | $3.67 \times 10^{12}$ |
| AAV5-CAG-GFP | $2.78 \times 10^{13}$ |
| AAV6-CAG-GFP | $4.35 \times 10^{12}$ |
| AAV8-CAG-GFP | $3.63 \times 10^{13}$ |
| AAV9-CAG-GFP | $2.37 \times 10^{13}$ |
| Spark100-CAG-GFP | $1.39 \times 10^{13}$ |
| Spark200-CAG-GFP | $2.09 \times 10^{13}$ |
| AAV1-Ef1a-Parkin (human) | $1.04 \times 10^{13}$ |
| AAV2-CAG-Parkin (human) | $2.43 \times 10^{13}$ |

**Table 3:** List of vectors used for in-vivo Studies.

| Name | Vector Titer (vg/mL) |
| --- | --- |
| Spark100-Ef1a- Parkin (human) | $3.56 \times 10^{13}$ |
| Spark100-CAG- Parkin (human) | $1.14 \times 10^{13}$ |
| Spark100-hSYN1- Parkin (human) | $7.92 \times 10^{13}$ |
| Spark100-TH (full)- Parkin (human) | $3.01 \times 10^{13}$ |
| Spark100-TH (mini)- Parkin (human) | $1.71 \times 10^{13}$ |
| AAV1-Ef1a-Parkin (human) | $1.04 \times 10^{13}$ |

### Protein Assays

#### Sample Processing

Dissected right SN samples were suspended in 500 uL radioimmunoprecipitation assay (RIPA) buffer (Thermo Fisher, Cat. No. PI89901) +PIC (Thermo Fisher, Cat. No. 78440) and homogenized using the TissueLyser II (QIAGEN, Cat. No. 85300) at 25.0 Hz for 30 seconds twice, switching sample arms between the repeats.

Total protein concentration in homogenized samples was determined with the BCA method using a commercial kit (Thermo Fisher, Cat. No. 23225) performed according to manufacturer’s instructions.

#### Capillary Electrophoresis

Protein lysate was diluted 1:5 in the provided sample buffer, then mixed with the master mix and heated at 75°C for 10 min. Protein mix containing 1 µg of protein, protein normalization reagent, blocking reagent, wash buffer, biotinylated ladder, primary antibodies (Table 4), and chemiluminescent substrate were dispensed into designated wells in a manufacturer-provided microplate. The plate was loaded into the instrument (Protein Simple, Cat #004-650.), and proteins were drawn into individual capillaries on a 25 capillary cassette (12–230 kDa). Data was analyzed using Compass software (v6.0 or higher) provided by the manufacturer. Protein concentrations were normalized to the total protein content in the capillary and reported as “corrected area under the curve” (A.U.C.). Values under 10,000 A.U.C. were considered below the level of quantitation (BLOQ).

#### pUb^Ser65^ ELISA

Assay was performed using the Cell Signaling PathScan pUb^Ser65^ Sandwich ELISA Kit (Cat#28384). An amount of protein lysate totaling 5ug of was added into Sample Diluent to a final volume of 100uL and loaded onto the plate. The plate was sealed and incubated at 37C for 2 hours with light shaking (250 RPM). Plate was washed 4 times with 200uL 1X wash buffer and wells were aspirated after the final wash. 100uL reconstituted antibody was added and the plate was again sealed and incubated at 37C for 1 hour. Previous wash steps and aspiration of wells were repeated. 100uL reconstituted HRP-linked secondary antibody was added, plate was sealed and incubated at 37 for 30 minutes. Wash step with aspiration was again repeated. 100uL of TMB Substrate was added and the plate was incubated in the dark at RT for 30 minutes. A 100uL stop solution was then added and plate was read on Biotek Synergy H1 Multimode Reader with Agilent Gen5 software. Absorbance was read at 450nm, and the standard curve was fit using a four-parameter logistic curve.

#### Immunostaining

Forty formaldehyde-perfused and fixed mouse brain hemispheres (one hemisphere per brain) were cryoprotected with 0.1 M phosphate buffer (PB, pH 7.4) containing 20% sucrose for 72 hours, and then rapidly frozen. Serial cryostat sections (30-µm) were cut coronally through the brainstem containing the substantia nigra (SN), approximately from Bregma −2.46 mm to −3.88 mm.

One set of sections was processed for double-immunofluorescence labeling (TH + Parkin). Following washes in 0.01 M phosphate buffered saline (PBS, pH 7.4), sections were incubated free-floating in PBS containing 0.3% Triton X-100, 2% blocking serum and 2 primary antibodies for 43 hours at 4°C. This was followed by incubation of sections in PBS containing 2 separate secondary antibodies conjugated with Alexa Fluor® 594 (for TH) and Alexa Fluor® 488 (for Parkin), respectively. Sections were then counterstained with FD DAPI solution™ (FD Neurotechnologies, Columbia, MD). All above steps were carried out at room temperature except where indicated, and each step was followed by washing in PBS. After thorough washes in PBS, all sections were mounted on gelatin-coated slides (4 sections per slide), cover slipped with Vectashield® (Vector Lab, Burlingame, CA), sealed with nail polish, and stored at 4°C.

A second set of sections was processed for triple-immunofluorescence labeling (TH + Iba1 + GFAP). All methods are identical to the above description with the exception of the secondary antibody incubation, which was performed in PBS containing 3 separate secondary antibodies conjugated with Alexa Fluor® 647 (for TH), Alexa Fluor® 594 (for Iba1) and Alexa Fluor® 488 (for GFAP).

Twenty-four PFA-perfused and fixed rat brain hemispheres (one hemisphere per brain) were cryoprotected with 0.1 M phosphate buffer (PB, pH 7.4) containing 20% sucrose for 72 hours, and then rapidly frozen. Serial cryostat sections (30-µm) were cut coronally containing the substantia nigra (SN). One set of sections was processed for double-immunofluorescence labeling (TH + Parkin following standard IHC procedure). After initial washes, sections were blocked with 5% BSA and 0.3% Triton X100 for 30 minutes and then incubated with primary antibody (please refer to the antibody list below) overnight at 4C. Sections were then incubated with a secondary antibody for 1.5 hours at room temperature. Washes with PBS were performed in between each step.

All imaging was performed on a Zeiss Axioscan Z1 using a 5x air objective at 0.91 μm per pixel.

#### HPLC Analysis of Catecholamines and Acidic Metabolites

Dopamine (DA), 3,4-dihydroxyphenylacetic acid (DOPAC), homovanillic acid (HVA) and serotonin (5-HT) concentrations in right striatal tissue samples were determined by high performance liquid chromatography (HPLC) with electrochemical detection.

The frozen tissue samples were homogenized (1:10 w/v) in a 0.1 M acetic acid, 1 mM oxalic acid, 3 mM L-cysteine - 0.2 M perchloric acid mixture (1:6) with ultrasonic processor (SONOPULS mini20 with MS 1.5 microtip, Bandelin Electronic GmbH & Co, Berlin, Germany). Tissue homogenates were centrifuged for 10 min at 17000 g at 4 °C. Supernatants were collected, centrifuged again, and transferred into glass vials for analysis.

The Infinity II 1260 HPLC system (Agilent Inc., Santa Clara, CA, USA) consists of a G711A solvent delivery system, G7129A thermostatic autosampler, amperometric electrochemical detector (Logichrom, Lablogic Systems Ltd., Sheffield, UK) equipped with Sencell GC 2mm flow cell, and a salt-bridge reference electrode. The HPLC system is controlled by the LAURA control and analysis software (LabLogic Systems Ltd.). The analytes were detected with a 0.78 V working electrode potential.

The analytes were separated on a Kinetex C-18 reversed-phase column, 4.6 ′ 100 mm, 2.6 µm, with a SecurityGuard Ultra precolumn for C18 columns (Phenomenex ApS, Værløse, Denmark) in an isocratic run. The column and electrochemical detector were maintained at 55 °C. The mobile phase is 100 mM phosphoric acid, 100 mM citric acid, 0.6 g/L 1-octanesulfonic acid, 12 mM NaCl and 0.1 mM disodium EDTA, pH 3.05 – acetonitrile mixture (1000:85, v/v). The flow rate was set to 1 mL/min and injection volume to 10 µL. The levels of the analytes were calculated using external standards and expressed as pg/g wet tissue.

**Table 7.** List of Primary Antibodies used in the study.

| Antigen |  | Host Species | Target Species | Tag | Dilution | Manufacturer | Cat. No. | Assay |
| --- | --- | --- | --- | --- | --- | --- | --- | --- |
| Primary Antibodies | GFAP | Rat | H, Ms, Rat | Unconjugated | 1:250 | ThermoFisher | 13-0300 | IHC |
|  | IBA1 | Rabbit | H, Ms, Rat | Unconjugated | 1:500 | FUJIFILM | 019-19741 | IHC |
|  | TH | Sheep | Mammalian | Unconjugated | 1:500 | Pel-Freez | P60101-150 | IHC |
|  | Parkin | Rabbit | H, Ms, Rat | Unconjugated | 1:1000 | Thermo Fisher | 702785 | IHC |
|  | GFAP | Rabbit | H, Ms, Rat | Unconjugated | 1:100 | Cell Signaling | 12389 | JESS |
|  | IBA1 | Rabbit | H, Ms, Rat | Unconjugated | 1:50 | Abcam | ab178847 | JESS |
|  | TH | Rabbit | Mammalia | Unconjugated | 1:100 | Pel-Freez | P40101-150 | JESS |
|  | Parkin | Mouse | H, Ms, Rat | Unconjugated | 1:100 | Cell Signaling | 4211 | JESS |
|  | Parkin | Rabbit | H, Ms, Rat | Unconjugated | 1:100 | Thermo Fisher | 702785 | JESS |
|  | Parkin | Mouse | H, Ms, Rat | Unconjugated | 1:250 | Santa-Cruz | sc-32282 | ICC fibroblast |
|  | pUbSer65 | Rabbit | H | Unconjugated | 1:250 | Cell Signaling | 62802 | ICC fibroblasts |
|  | pUbSer65 | Rabbit | H | Unconjugated | 1:500 | Cell Signaling | 62802 | Dot Blot |

GFAP = glial fibrillary acidic protein; H = human; IBA1 = ionized calcium-binding adapter

#### Statistical analysis

All quantitative data in the figures, unless otherwise specified, are presented as mean ± SD and analyzed using GraphPad Prism software (version 10.3.0) or MS Excel. Comparisons between the two groups were done using unpaired two-tailed Student’s t test. Differences between more than two groups were carried out using 1-way ANOVA with Tukey’s or Dunnett’s correction for relevant pairwise comparisons. For all statistical analysis, *p* < 0.05 was considered significant.

The statistical analysis performed for each dataset is indicated in the figure legends.

#### Sequence of Promoters

##### CAG

ATCTAGTTATTAATAGTAATCAATTACGGGGTCATTAGTTCATAGCCCATATATGGAGTTCCGCGTTACATAACTTACGGTAAATGGCCCGCCTGGCTGACCGCCCAACGACCCCCGCCCATTGACGTCAATAATGACGTATGTTCCCATAGTAACGCCAATAGGGACTTTCCATTGACGTCAATGGGTGGAGTATTTACGGTAAACTGCCCACTTGGCAGTACATCAAGTGTATCATATGCCAAGTACGCCCCCTATTGACGTCAATGACGGTAAATGGCCCGCCTGGCATTATGCCCAGTACATGACCTTATGGGACTTTCCTACTTGGCAGTACATCTACGTATTAGTCATCGCTATTACCATGGTCGAGGTGAGCCCCACGTTCTGCTTCACTCTCCCCATCTCCCCCCCCTCCCCACCCCCAATTTTGTATTTATTTATTTTTTAATTATTTTGTGCAGCGATGGGGGCGGGGGGGGGGGGGGGGCGCGCGCCAGGCGGGGCGGGGCGGG GCGAGGGGCGGGGCGGGGCGAGGCGGAGAGGTGCGGCGGCAGCCAATCAGAGCGGCGCGCTCCGAAAGTTTCCTTTTATGGCGAGGCGGCGGCGGCGGCGGCCCTATAAAAAGCGAAGCGCGCGGCGGGCGGGAGTCGCTGCGCGCTGCCTTCGCCCCGTGCCCCGCTCCGCCGCCGCCTCGCGCCGCCCGCCCCGGCTCTGACTGACCGCGTTACTCCCACAGGTGAGCGGGCGGGACGGCCCTTCTCCTCCGGGCTGTAATTAGCGCTTGGTTTAATGACGGCTTGTTTCTTTTCTGTGGCTGCGTGAAAGCCTTGAGGGGCTCCGGGAGGGCCCTTTGTGCGGGGGGAGCGGCTCGGGGGGTGCGTGCGTGTGTGTGTGCGTGGGGAGCGCCGCGTGCGGCTCCGCGCTGCCCGGCGGCTGTGAGCGCTGCGGGCGCGGCGCGGGGCTTTGTGCGCTCCGCAGTGTGCGCGAGGGGAGCGCGGCCGGGGGCGGTGCCCCGCGGTGCGGGGGGGGCTGCGAGGGGAACAAAGGCTGCGTGCGGGGTGTGTGCGTGGGGGGGTGAGCAGGGGGTGTGGGCGCGTCGGTCGGGCTGCAACCCCCCCTGCACCCCCCTCCCCGAGTTGCTGAGCACGGCCCGGCTTCGGGTGCGGGGCTCCGTACGGGGCGTGGCGCGGGGCTCGCCGTGCCGGGCGGGGGGTGGCGGCAGGTGGGGGTGCCGGGCGGGGCGGGGCCGCCTCGGGCCGGGGAGGGCTCGGGGGAGGGGCGCGGCGGCCCCCGGAGCGCCGGCGGCTGTCGAGGCGCGGCGAGCCGCAGCCATTGCCTTTTATGGTAATCGTGCGAGAGGGCGCAGGGACTTCCTTTGTCCCAAATCTGTGCGGAGCCGAAATCTGGGAGGCGCCGCCGCACCCCCTCTAGCGGGCGCGGGGCGAAGCGGTGCGGCGCCGGCAGGAAGGAAATGGGCGGGGAGGGCCTTCGTGCGTCGCCGCGCCGCCGTCCCCTTCTCCCTCTCCAGCCTCGGGGCTGTCCGCGGGGGGACGGCTGCCTTCGGGGGGGACGGGGCAGGGCGGGGTTCGGCTTCTGGCGTGTGACCGGCGGCTCTAGAGCCTCTGCTAACCATGTTCATGCCTTCTTCTTTTTCCTACAGCTCCTGGGCAACGTGCTGGTTATTGTGCTGTCTCATCATTTTGGCAAAGAATT

##### EF1a

GGGCAGAGCGCACATCGCCCACAGTCCCCGAGAAGTTGGGGGGAGGGGTCGGCAATTGAACCGGTGCCTAGAGAAGGTGGCGCGGGGTAAACTGGGAAAGTGATGTCGTGTACTGGCTCCGCCTTTTTCCCGAGGGTGGGGGAGAACCGTATATAAGTGCAGTAGTCGCCGTGAACGTTCTTTTTCGCAACGGGTTTGCCGCCAGAACACAG

##### hSYN11

AGTGCAAGTGGGTTTTAGGACCAGGATGAGGCGGGGTGGGGGTGCCTACCTGACGACCGACCCCGACCCACTGGACAAGCACCCAACCCCCATTCCCCAAATTGCGCATCCCCTATCAGAGAGGGGGAGGGGAAACAGGATGCGGCGAGGCGCGTGCGCACTGCCAGCTTCAGCACCGCGGACAGTGCCTTCGCCCCCGCCTGGCGGCGCGCGCCACCGCCGCCTCAGCACTGAAGGCGCGCTGACGTCACTCGCCGGTCCCCCGCAAACTCCCCTTCCCGGCCACCTTGGTCGCGTCCGCGCCGCCGCCGGCCCAGCCGGACCGCACCACGCGAGGCGCGAGATAGGGGGGCACGGGCGCGACCATCTGCGCTGCGGCGCCGGCGACTCAGCGCTGCCTCAGTCTGCGGTGGGCAGCGGAGGAGTCGTGTCGTGCCTGAGAGCGCAG

##### TH (Full)

CTGCCTCCAGAGAGCCTGGCCCCAAGGAAGAGTCTAGTAAGCTTAGTTCCCATCGGGCTTCCATGAAAGCACAACTGGCCCGGCAGGAAACCGAATTAAAAAGCAATATTTGTATCAGTGGAAGACATTTGCTGAAAGGTTAAATCCACATCCGGCAGTGTGGGCCATGAGCCTCCGGCGTGGTGTTCATCAGGCATGTCTCTCCTCCTGGCCTGGGCACCTGAGCACTGGGGCTGCCCTGGGCAGAGCTGGGGCAGGGTGCTGGGGGGCCTGGAGCTGCCTCACCGAGGGATCCTCAGCAGCCGACCCTGGGGGAGGCAAATGAGACTCTTTCTGGGG ACCTTGAGGGGAGCTCGGGGGAGCCATGCAGAGCTTCACCAGGCCTGGACACTGGGCATGGAGGCTGGGCCACCCAAGGGCCATCACCAGGGACTCAGGTGGGTGGGCCTCAGCCCTGGGTGACAGAAGCTCACGGGCTGCAGGGCGAGGCCAGAGGCTGAGCCTTCAGGCTGAGGTCTTGGAGGCAAATCCCTCCAACGCCCTTCTGAGCAGGCACCCAGACCTACTGTGGGCAGGACCCACAGGAGGTGGAGGCCTTTGGGGAACACCGTGGAGGGGCATAGCATCTCCGAGAGAGGACAGGGTCTGCACTGGGTGCTGAGAGACAGCAGGGGCCGAGCGGTAGGCTTCCCTGCCCCCAGGGATGTTCCAGGGGAGCGCAAGGGAGGGGCATTAATATCGTGGCAAGAAAGGGCAGGCATTGCAGAGTGAGCAGCGACGGAACTGGGTTTTGTGGGATGCATAGGAGTTCACCCGGATAAGAGGTGGGTGAGGAATGACACTGCAAACCGGGGATCACGGAGCCCCAAATCCTTCTGGGCCAGGAAGTGGGAAGGGTTGGGGGGTCTTCCCTTTGCTTTGACTGAGCACTCAGCCTGCCTGCAGAGGGCAGCGAGGAGCCACGGAGGGGTGTGGGACAGGGATGCCATGGCTGAAGCAGTTTTAGGAAAGGTCCCAGGGGCTATTGTTGAAGAGAGAACGGGGAGCGGGGAGTCCCACAGCTGACAGGAGCAGAGTGGGCCCTGAGAGATGCCAGCTCTGGGTGCCACAGTGACCAGCCGGGGTAGGCCTTCGAGAAGTCAGGGAGCGTCTAGGGCTTCTGGCTCCTGCTGGGCCCAGGGTGTCATCTTGGGCTGCCAACACCAGAAAGCCCAGCAGATACAGGAAGCCCCAAGCCCTGTCGGAAACGGTTCTTCTCCAGGAGGGACAGCGGTGGCAGCGTTCAGCCGCAGGCCATGCACTCTGGGGCCACGTCCTTCCCTCTGTACAGTCCAGCATTGTCAAGGCGGGCTCTGGCCATCTCTGCTGACCCCAGAGGGATGGGGAGGCCTCCCCTTCCACCAGAAGGGCCAGAAGCCACCCTGGGCAGGGGCATCACTCTCCCTGGGTGGGGCAGCGGCGGGGAGCAGGAGGTGCCAGTGGGCGTGGGCTGGATGCGGGTGCCTGCGGGGCGGACATGGAACTTGGGGGAGGCTCTAGGCTGGGGTTGTCCTCAAGGGAGTTCTCAGGTCACCCCAGGGTCACCCTCAACCCGGGGCCTGGTGGGGTAGAGGAGAAACTGCAAAGGTCTCTCCAAGGGGAAGGCATCAGGGCCCTCAGCACTGAGGGACGTGCGTGCTCTTCAAAGAAGGGGCCACAGGACCCCGAGGGAAGCCAGGAGCTAGCAGTGGGCCATAGAGGGGCTGAGTGGGGTGGGTGGAAGCCGTCCCTGGCCCTGGTCGCCCTGGCAACCCTGGTGGGGACTGTGATGCAGGAGGTGGCAGCCATTTGGAAACGCGTGGCGTCTCCTTAGAGATGTCTTCTTCAGCCTCCCAGGGTCCTCCACACTGGACAGGTGGGCCCTCCTGGGACATTCTGGACCCCACAGGGCGAGCTTGGGAAGCCGCTGCAAGGGCCACACCTGCAGGGCCCGGGGGCTGTGGGCAGATGGCACTCCTAGGAACCACGTCTACAAGACACACGGCCTGGAATCTTCTGGAGAAGCAAACAAATTGCCTCCTGACATCTGAGGCTGGAGGCTGGATTCCCCGTCTTGGGGCTTTCTGGGTCGGTCTGCCACGAGGTTCTGGTGTTCATTAAAAGTGTGCCCCTGGGCTGCCAGAAAGCCCCTCCCTGTGTGCTCTCTTGAGGGCTGTGGGGCCAAGGGGACCCTGGCTGTCTCAGCCCCCCGCAGAGCACGAGCCCCTGGTCCCCGCAAGCCCGCGGGCTGAGGATGATTCAGACAGGGCTGGGGAGTGAAGGCAATTAGATTCCACGGACGAGCCCTTTCTCCTGCGCCTCCCTCCTTCCTCACCCACCCCCGCCTCCATCAGGCACAGCAGGCAGGGGTGGGGGATGTAAGGAGGGGAAGGTGGGGGACCCAGAGGGGGCTTTGACGTCAGCTCAGCTTATAAGAGGCTGCTGGGCCAGGGCTGTGGAGACGGAGCCCGGACCTCCACACTGAGCCA

##### TH (mini)

AGACACACGGCCTGGAATCTTCTGGAGAAGCAAACAAATTGCCTCCTGACATCTGAGGCTGGAGGCTGGATTCCCCGTCTTGGGGCTTTCTGGGTCGGTCTGCCACGAGGTTCTGGTGTTCATTAAAAGTGTGCCCCTGGGCTGCCAGAAAGCCCCTCCCTGTGTGCTCTCTTGAGGGCTGTGGGGCCAAGGGGACCCTGGCTGTCTCAGCCCCCCGCAGAGCACGAGCCCCTGGTCCCCGCAAGCCCGCGGGCTGAGGATGATTCAGACAGGGCTGGGGAGTGAAGGCAATTAGATTCCACGGACGAGCCCTTTCTCCTGCGCCTCCCTCCTTCCTCACCCACCCCCGCCTCCATCAGGCACAGCAGGCAGGGGTGGGGGATGTAAGGAGGGGAAGGTGGGGGACCCAGAGGGGGCTTTGACGTCAGCTCAGCTTATAAGAGGCTGCTGGGCCAGGGCTGTGGAGACGGAGCCCGGACCTCCACACTGAGCCATGCCCACCCCCGACGCC ACCACGCCACAGG

